# The vitals for steady nucleation maps of spontaneous spiking coherence in autonomous two-dimensional neuronal networks

**DOI:** 10.1101/2024.01.05.574396

**Authors:** Dmitrii Zendrikov, Alexander Paraskevov

## Abstract

Thin pancake-like neuronal networks cultured on top of a planar microelectrode array have been extensively tried out in neuroengineering, as a substrate for the mobile robot’s control unit, i.e., as a cyborg’s brain. Most of these attempts failed due to intricate self-organizing dynamics in the neuronal systems. In particular, the networks may exhibit an emergent spatial map of steady nucleation sites (“n-sites”) of spontaneous population spikes. Being unpredictable and independent of the surface electrode locations, the n-sites drastically change local ability of the network to generate spikes. Here, using a spiking neuronal network model with generative spatially-embedded connectome, we systematically show in simulations that the number, location, and relative activity of spontaneously formed n-sites (“the vitals”) crucially depend on the samplings of three distributions: 1) the network distribution of neuronal excitability, 2) the distribution of connections between neurons of the network, and 3) the distribution of maximal amplitudes of a single synaptic current pulse. Moreover, blocking the dynamics of a small fraction (about 4%) of non-pacemaker neurons having the highest excitability was enough to completely suppress the occurrence of population spikes and their n-sites. This key result is explained theoretically. Remarkably, the n-sites occur taking into account only short-term synaptic plasticity, i.e., without a Hebbian-type plasticity. As the spiking network model used in this study is strictly deterministic, all simulation results can be accurately reproduced. The model, which has already demonstrated a very high richness-to-complexity ratio, can also be directly extended into the three-dimensional case, e.g., for targeting peculiarities of spiking dynamics in cerebral (or brain) organoids. We recommend the model as an excellent illustrative tool for teaching network-level computational neuroscience, complementing a few benchmark models.

## 1. Introduction

Spatiotemporal patterns are the main source of information about the surrounding world: each phenomenon or event is characterized by where and when it happens. In the neuroscience domain, it has been recently shown that unique spatiotemporal patterns of functional connections in the human brain can be used, like fingerprints, to identify a particular person [1–3]. However, studies of such an identification require expensive scanners for functional magnetic resonance imaging or magnetoencephalography, while invasive experimental techniques are limited due to ethical considerations. Besides, the human brain itself is so complex that it is extremely difficult to reliably distinguish a true statistically significant effect from a multitude of artifacts.

Therefore, it seems worth turning first to simpler “toy” experimental systems, in particular, to two-dimensional (2D) neuronal networks grown *in vitro*, like a woven carpet, the so-called neuronal cultures [4, 5]. Dense micro-electrode arrays embedded in the surface under such a carpet allow detecting electrical impulse (“spiking”) activity of neurons with high spatial and temporal resolution. It is well-known that neuronal cultures demonstrate a rich repertoire of characteristic spatiotemporal patterns of spontaneous spiking activity in the form of the so-called population spikes (PSs) or bursts of such spikes [6–14]. Interestingly, PSs occur from nucleation sites (n-sites) and each mature neuronal culture usually has several n-sites [15–20]. The specific number of n-sites and their location (i.e., the spatial map of n-sites) are likely unique for each network, serving as reliable natural marks of distinction of the networks from each other. Despite the comparative simplicity and reproducibility of the *in vitro* experiments, which are also qualitatively reproduced in simulations by spiking neuronal network models [19–22], a detailed mechanism underlying the occurrence of steady spatial map of n-sites for spontaneous PSs remains elusive. Meanwhile, similar phenomena typically associated with the so-called neuronal avalanches subject to self-organized criticality (SOC) can also occur *in vivo* [23–25] (on SOC in brain slices and neuronal cultures *in vitro*, see [26, 27] and [28–30]). In addition, as population spikes originating from a steady n-site mimic events of focal epilepsy, spontaneously formed n-sites might be used as a simplistic prototype of focal epilepsy foci on the cortical sheet [31, 32]. So understanding the mechanism of forming the map of n-sites seems quite important.

In its turn, such an understanding arises as the result of constructing a theory that consistently explains key statistically-proven dependencies. However, a weeks-long period of neuronal culture growth, difficulties in getting structurally similar samples with the spatial homogeneity of the location of neurons on the substrate, and other restricting factors make it very challenging to obtain reliable experimental statistics for the spatial maps of n-sites. Fortunately, as mentioned above, there are mathematical models of neuronal cultures that plausibly simulate spatiotemporal patterns of neuronal activity observed in the “wet” experiments: the models describe an occurrence of episodic PSs from a small number of spontaneously-formed steady n-sites [19–22].

In this article, we present the results of systematic simulations performed to study the key parametric dependencies of the 2D spiking neuronal network model [22] that affect stationary spatial map of n-sites. An important feature of this generative model is that it is completely deterministic, i.e., all of its results are accurately reproducible. To identify the influence of the model parameters on the occurring nucleation maps, we initially prepared a certain reference realization of the neuronal network, and obtained the reference nucleation map for it. Further, by changing the target parameters of the reference neuronal network (RefNN), we observed how the nucleation pattern changed in response.

As a result, we have found that the number, location, and relative activity of PS n-sites crucially depend on specific samplings of three distributions: 1) the network distribution of neuronal excitability, 2) the distribution of connections between neurons of the network, and 3) the distribution of maximal amplitudes of the synaptic current pulse.

Aside from the apparent use for advanced computational modeling of focal epilepsy [33– 35], these results indicate that the nucleation pattern can change significantly (i) with the neuronal network age (cp. [36]) and (ii) due to external electrical stimulation, especially spatially local one, when only a part of the network is stimulated [37–41]. The latter is standardly used in attempts to create a hybrid bio-electronic control system for a mobile robot (e.g., [42–44], reviewed in [45, 46]).

From the viewpoint of physics, both planar neuronal cultures and the suggested neuronal network model in particular exhibit a bright example of autowave pattern-forming system out of equilibrium [47–49], similarly to the family of Belousov-Zhabotinsky reactions [50, 51]. For instance, it has been previously shown in simulations with the model that in relatively dense networks, where inhibitory neurons are not blocked, a PS can originate in the form of multi-armed spiral wave with the drifting center [22]. If connections between neurons in the model network are rather sparse and the inhibitory neurons are blocked, as we consider here, then a PS occurs only in the form of outward circular waves beginning in a few steady n-sites, which originate spontaneously. As in many other pattern-forming dissipative systems, the nature of such a spontaneous nucleation is insufficiently clear. The previous study has shown that n-sites are not located at local inhomogeneities of the spatial density of neurons, which is on average homogeneous [22]. In the present study, we have found that the nucleation is crucially dependent on a small fraction of non-pacemaker neurons having the highest subcritical excitability: blocking the dynamics of these neurons leads to a complete suppression of PSs and their n-sites.

## 2. Model, its Analysis, and Methods

### 2.1. Neuronal network model

It consists of three main components: (i) an algorithm for generating the network connectome, (ii) a spiking neuron model, and (iii) a dynamic synapse model describing the interaction between neurons. The network standardly has 80% excitatory and 20% inhibitory neurons [52–56]. However, in order to obtain a clear map of n-sites, the inhibitory neurons have been blocked (i.e., their membrane potentials have been clamped to the resting potential *V*_*rest*_) in most of the simulations described in this article. The impact of inhibitory neurons is described in **Discussion**; Suppl. Video 2 shows a simulation with unblocked inhibitory neurons (see also [57]).

#### 2.1.1. Network connectome model

We assume that (i) *N* point neurons are uniformly distributed over square area *L× L* and (ii) the probability of formation of a unilateral connection between each pair of neurons decreases exponentially as a function of the distance *r* between them, if the distance is not too large [58–66],

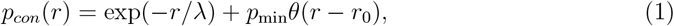

where 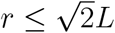, λ is the characteristic connection length (both *r* and λ are further expressed in units of *L* by default), *r*_0_ = λ ln(1/*p*_*min*_) (see the description below), and *θ*(…) is the unit step function: *θ*(*x*) = 1 for *x* > 0 and *θ*(*x*) = 0 for *x* ≤ 0. In engineering, the exponential dependence *p*_*con*_(*r*) ∝ exp(−*r*) is sometimes referred to as the Waxman model [67, 68].

The second summand in Eq. (1) contributes to formation of long-range connections, typically increasing the total number of connections up to 5% (see the left graph in Fig. 1). This summand must be included to take into account constraints of the random number generator (RNG), which is necessary for generating the network connectome. In particular, the standard RNG in C can generate a random integer number within the range from 0 to some predetermined value *RAND MAX*, which may depend on the operating system but cannot be smaller than 32767. That integer number can be then transformed into the real one by normalizing it by *RAND MAX*, resulting in the pseudo-uniform distribution from 0 to 1 that is used to generate neuronal connections by comparing the random numbers with the probability *p*_*con*_. However, as the smallest nonzero random number is *p*_*min*_ = 1/*RAND MAX*, there is a systematic bias for nonzero probability values smaller than this limit: these values are compared with zero and always lead to the connection formation. If *RAND MAX* turns to infinity, the second term in Eq. (1) vanishes and we return to the Waxman model (i.e., the first summand in Eq. (1)). Given *RAND MAX* = 32767 and λ = 0.01, one gets *p*_*min*_ ≈ 3 *·* 10^−5^ and *r*_0_ ≈ 0.1. A comparative study of neuronal networks generated with and without the second summand in Eq. (1) is given in Ref. [69]. In brief, for *N* = 50000 and λ = 0.01 standardly used in this study, the average number of outgoing connections per neuron increases from 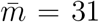 for the Waxman model to 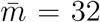 for Eq. (1), with the unchanged standard deviation (= 6) (cp. [70]). Thus, effectively, every neuron acquires one additional long-range connection [71]. As the fraction of additional connections is quite small, the results of comparative simulations have been qualitatively the same.

**FIG. 1.**
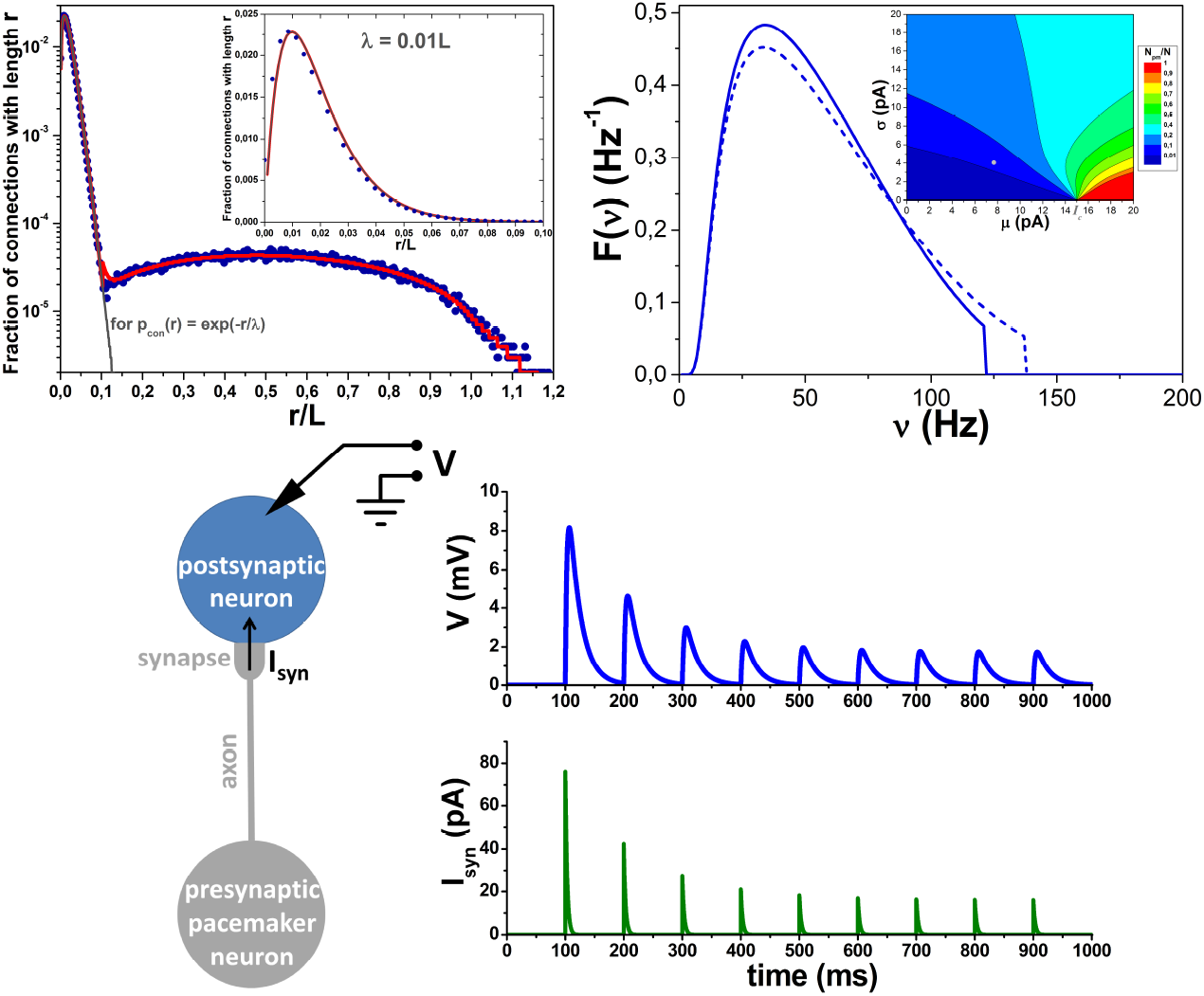
Upper graphs: LEFT: The red curve is a fraction of network connections with the given length *r*, or the product *p*_*con*_(*r*)*P* (*r*), where *p*_*con*_(*r*) and *P* (*r*) are given by Eqs. (1) and (3), respectively. In turn, the gray curve illustrates the case *p*_*con*_(*r*) = exp(−*r/λ*) following from Eq. (1) at *p*_min_ = 0. The blue dots represent the data for the RefNN network connectome used in all simulations. The inset shows the same graph with linear scale on the vertical axis and *r/L* ≤ 0.1. RIGHT: Probability density function *F* (*ν*), given by Eq. (12), for self-frequencies *ν* of pacemaker neurons (solid line is for excitatory neurons, dashed line is for inhibitory neurons; the difference originates from the different values of *τ*_*ref*_ ), if the background currents are distributed by the non-negative and upper-bounded normal distribution, see Eq. (9). Inset: Fraction of pacemaker neurons *N*_*pm*_*/N* (formula (13) with *I*_max_ = 20 pA) as a function of two basic parameters for the normal distribution (9) of background currents the mean *µ* and standard deviation *σ*. The filled gray circle indicates the values used in simulations. The critical value of the background current, above which the neuron is a pacemaker, is *I*_*c*_ = 15 pA. At the bottom: LEFT: Schematic outline of two LIF neurons, see Eq. (7), coupled by a monosynaptic excitatory connection. The presynaptic neuron is a newly minted pacemaker (with a spiking frequency of 10 Hz), while the postsynaptic one is not. The synaptic current *I*_*syn*_(*t*) = *Jy*(*t*), where *t* is time, is subject to a short-term depression of fraction *y*(*t*) determined by Eqs. (15). RIGHT: The time dependencies for the postsynaptic neuron potential *V* counted from the resting potential (top) and *I*_*syn*_ (bottom). The constant *J* was taken of 152 pA, the maximal value in the samplings for all simulations. Provided that spiking threshold potential *V*_*th*_ = 15 mV, the dynamics show that even the largest (not depressed) first pulse of *I*_*syn*_ is insufficient to cause a spike generation on the postsynaptic neuron if the latter is not fed by non-zero subcritical background current.

It is important to stress that, regardless of the second summand in Eq. (1) for *p*_*con*_(*r*), the considered networks belong to the small-world network type [72–76], with the following mean values of the clustering coefficient (CC) and the shortest path length (SPL): CC ≈ 0.13 and SPL ≈ 4 for Eq. (1), and CC ≈ 0.15 and SPL ≈ 11 for the Waxman model [69]. Based on these numbers, the “small-world-ness” introduced in [73] is increased more than twice (from about 45 to 103) with including the second summand in Eq. (1). Therefore, the RNG-caused correction in Eq. (1) substantially *improves* this criterion.

It is also worth noting that inequality 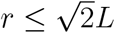 implies that connections between neurons are geometrically modeled by segments of straight lines [77–79]. In other words, we assume that the connection length is simply equal to the distance between neurons. As the square area is a convex set of points, this simplification seems rational. We emphasize that the connections do not cross boundaries of the square and, therefore, the neurons in the vicinity of the boundaries have fewer connections. For simplicity, parameter λ in Eq. (1) is chosen independently of whether the preand postsynaptic neurons are excitatory or inhibitory (cp. [59]). The formation of autaptic connections (i.e., self-connections) [80] is also prohibited to this end (cp. [81]). Duplicate connections are not considered in the model as well: every neuron can have only one outgoing synapse to any other neuron.

With a given function *p*_*con*_(*r*), the average number of connections in a network of *N* neurons is

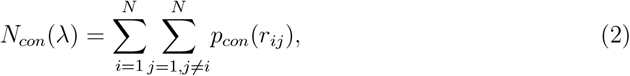

where *r*_*ij*_ is the distance between the *i*-th and *j*-th neurons. For the neurons being uniformly distributed over the square *L × L, r*_*ij*_ is a random value with the distribution density given by (*r* is expressed in units of *L*)

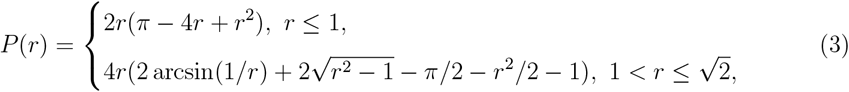

Here P(*r*) is the probability density to find two point neurons at distance *r* from each other, such that 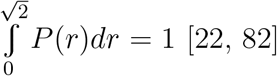. It has a single maximum 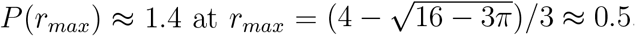.

The resulting distribution of connection lengths is given by the product *p*_*con*_(*r*)*P*(*r*), which, as a function of *r*, reaches its maximum at *r* ≈ λ for λ ≲ 0.1 and *r < r*_0_ (see Fig. 1, the inset of the left graph). The average number of connections in the network of *N* neurons is 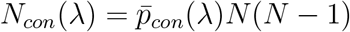, where the space-averaged probability

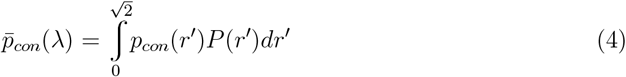

increases monotonically with λ, asymptotically reaching unity at infinity, and at λ ≪ 1 it has quite simple form 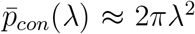. In turn, the average number of outgoing (or incoming) connections per neuron is 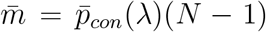. The average fraction *n*_*con*_(*r*) of network connections with the lengths longer than or equal to *r*, where 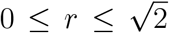, can also be determined explicitly (see Fig. S1 in Suppl. Mat.),

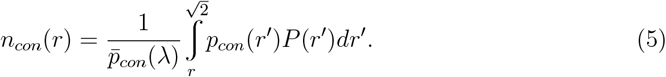

Finally, due to the fact that every connection between neurons has a metric length, there exist spike propagation delays. Assuming a constant speed of spike propagation along connections (implying axons), the delays are calculated by formula

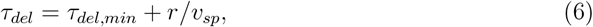

where *τ*_*del*_ is the total spike-propagation delay for a connection of length *r, τ*_*del*,*min*_ is the minimal delay same for all connections, and *v*_*sp*_ is the constant speed of spike propagation along the axon. We set *τ*_*del*,*min*_ = 0.2 ms and *v*_*sp*_ = 0.2 *L*/ms [83] with *L* = 1 mm by default.

#### 2.1.2. Neuron model

We use the standard Leaky Integrate-and-Fire (LIF) neuron that has no ability for intrinsic bursting. Subthreshold dynamics of transmembrane potential *V* of such a neuron is described by equation

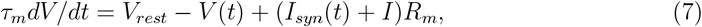

where *V*_*rest*_ is the neuron’s resting potential, *τ*_*m*_ is the characteristic time for relaxation of

*V* to *V*_*rest*_, *R*_*m*_ is the electrical resistance of the neuron’s membrane, *I*_*syn*_(*t*) is the total incoming synaptic current, which, as a function of time *t*, depends on the dynamic model of a synapse and the number of incoming synapses, *I* is a constant “background” current, the magnitude of which varies from neuron to neuron. The background currents determine the diversity of neuronal excitability and the fraction of pacemaker neurons in the network.

When the transmembrane potential reaches a threshold value *V*_*th*_ = *V* (*t*_*sp*_), it is supposed that the neuron emits a spike: *V* abruptly drops to a specified value *V*_*reset*_, *V*_*rest*_ ≤ *V*_*reset*_ *< V*_*th*_, and retains this value during the absolute refractory period *τ*_*ref*_ , after which the potential dynamics is again described by Eq. (7). The outcome of the LIF neuron dynamics for the rest of the network is a sequence of spike generation moments 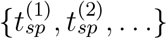.

If the LIF neuron has *I* value that exceeds *I*_*c*_ = (*V*_*th*_ −*V*_*rest*_)/*R*_*m*_, the critical (or rheobase) value, then this neuron is a pacemaker, i.e., it is able to emit spikes periodically with frequency

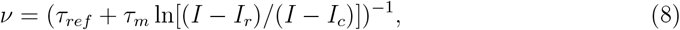

where *I*_*r*_ = (*V*_*reset*_ − *V*_*rest*_)/*R*_*m*_, in the absence of incoming signals from other neurons. Based on experimental findings [84, 85], we assume that both excitatory and inhibitory neurons may be pacemakers. In turn, if the background current *I* is less than *I*_*c*_, then this leads to an increase of depolarization of the neuron’s potential to some asymptotic subthreshold value, i.e. to the effective individualization of the neuronal resting potential, 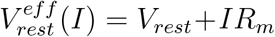.

In what follows, we consider that the background current values are distributed according to the non-negative and upper-bounded part of the normal (Gaussian) distribution, with the mean *µ* and standard deviation *σ*,

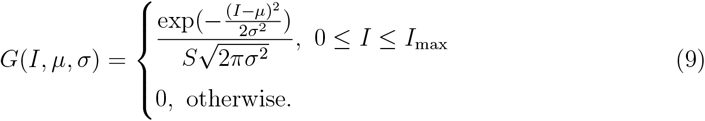

Here, *I*_*max*_ is the upper value of the background current and 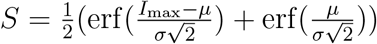, where erf(…) represents the error function, is a normalization factor ensuring equality 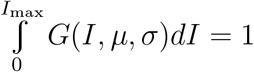.

In turn, the distribution density for self-frequencies (8) of the pacemaker neurons is given by formula (cp. [86])

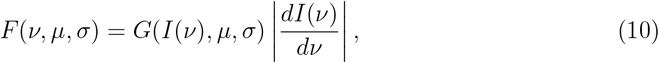

where

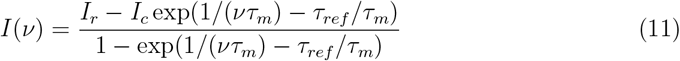

is the inverse function for *ν*(*I*), see Eq. (8). Explicitly, one gets (dependencies on *µ* and *σ* are implied)

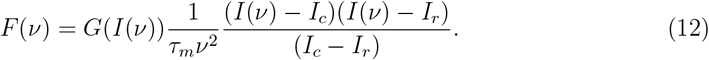

The distribution density *F* (*ν*) is plotted in Fig. 1 (right graph). The number of pacemakers *N*_*pm*_ can also be derived explicitly

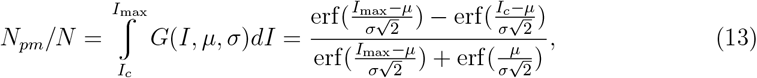

The corresponding plot for *N*_*pm*_/*N* given by Eq. (13) is shown on the inset of the right graph in Fig. 1. In all simulations we assume such inequalities *σ<µ<I*_*c*_ that *N*_*pm*_ ≪ *N*. Numerical values of parameters for the LIF neuron model: *τ*_*m*_ = 20 ms, *R*_*m*_ = 1 GΩ, *V*_*rest*_ = 0 mV, *V*_*th*_ = 15 mV, *V*_*reset*_ = 13.5 mV. These give the critical current value *I*_*c*_ = 15 pA and *I*_*r*_ = 13.5 pA. Refractory period *τ*_*ref*_ = 3 ms for excitatory neurons, *τ*_*ref*_ = 2 ms for inhibitory neurons. The non-negative part of normal distribution for the background currents, bounded above by *I*_ma*x*_ = 20 pA, has the mean *µ* = 7.7 pA and the standard deviation *σ* = 4.0 pA. These give the fraction (13) of pacemakers *N*_*pm*_/*N* = 3.4% with the maximal *ν* value 121 Hz for excitatory neurons and 138 Hz for inhibitory ones (see the right graph in Fig. 1).

#### 2.1.3. Synapse model

A single contribution to the incoming synaptic current in our neuronal network model is determined as

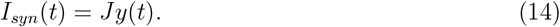

Here *J* is a constant that determines the amplitude of synaptic current impulse. The sign and magnitude of *J* depend on the type of preand postsynaptic neurons (i.e., whether the neuron is excitatory or inhibitory). Next, *y*(*t*) is a dimensionless parameter, 0 ≤ y ≤ 1, the dynamics of which is determined by the following system of equations [87, 88]:

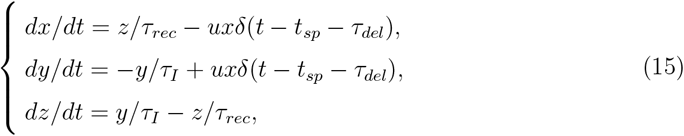

where *x, y* and *z* are the fractions of synaptic resources in the recovered, active and inactive states, respectively; *x* + y + *z* = 1; *τ*_*rec*_, *τ*_*I*_ are the characteristic relaxation times, *δ*(…) is the Dirac delta function, *t*_*sp*_ is the moment of spike generation at the presynaptic neuron, *τ*_*del*_ is the spike propagation delay (see Eq. (6)), and u is the fraction of recovered synaptic resource used to transmit the signal across the synapse, 0 ≤ *u* ≤ 1. The stable fixed point of Eqs. (15) is (*x, y, z*) = (1, 0, 0). The bottom plots in Fig. 1 show typical dynamics resulting from Eqs. (15) for a synapse between two excitatory neurons.

For the outgoing synapses of inhibitory neurons, the dynamics of u is described by equation [87, 88]

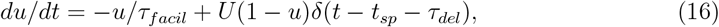

where *τ*_*facil*_ is the characteristic relaxation time, and 0 *<U* ≤ 1 is a constant parameter. For the outgoing synapses of excitatory neurons, *u* remains constant and equals to U.

Qualitatively, Eqs. (15) describe a short-term synaptic depression due to the depletion of synaptic resources at a high enough frequency of incoming spikes, while the additional Eq. (16) enables inhibitory synapses to overcome the depression and even increase their impact in a certain frequency range [87]. Focusing on excitatory neurons and synaptic depression, let us outline an important case of an outgoing synapse of the pacemaker neuron generating spikes with frequency *ν* given by Eq. (8). One can find the dependence of a stationary amplitude of synaptic current pulses on *ν* by simplifying Eqs. (15) as follows. Assuming that *τ*_*I*_ ≪ *τ*_*rec*_ and completely neglecting the finite-time dynamics of y (see the bottom right graph in Fig. 1), equations for the amplitudes (denoted by subscript “a”) of the fractions of synaptic resources read: *z*_*a*_ ≈ 1 − *x*_*a*_, *y*_*a*_ ≈ *ux*_*a*_, and *z*_*a*_/*τ*_*rec*_ − u*x*_*a*_*ν* = 0. Provided that *u* = *U*, these result in

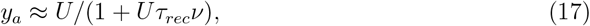

so the amplitude of synaptic current, see Eq. (14), is *Jy*_*a*_ ∼ 1/*ν* for relatively large *ν* values [89, 90]. This hyperbolic decrease is a direct consequence of the synaptic depression that may substantially reduce the impact of excitatory pacemakers on the other neurons of the network. A possibility of collective synchronization in excitatory spiking networks without synaptic depression has been considered, e.g., in [91]. To gain a qualitative understanding of the impact of short-term synaptic depression on network activity, one can recommend the mean-field analysis suggested in [92].

In the numerical simulations all synaptic parameters, except *τ*_*I*_, were normally distributed with the mean values *µ*_*k*_ described below, i.e., each synapse had its own unique values of these parameters. Standard deviations for all distributed parameters were equal to 0.5*µ*_*k*_. The maximal (or minimal, if the parameter is negative) values of the distributions were restricted by 4*µ*_*k*_ (or 1, if 4*µ*_*k*_ *>* 1 for parameter U), and the minimal (maximal, if the parameter is negative) values were restricted by zero (or the time step, for the time constants).

The numerical values of parameters for the synapse model were as follows [88]: *τ*_*I*_ = 3 ms, mean values for the normal distributions *τ*_*rec*,*ee*_ = *τ*_*rec*,*ei*_ = 800 ms, *τ*_*rec*,*ie*_ = *τ*_*rec*,*ii*_ = 100 ms, *τ*_*facil*,*ie*_ = *τ*_*facil*,*ii*_ = 1000 ms, *J*_*ee*_ = 38 pA, *J*_*ei*_ = 54 pA, *J*_*ie*_ = *J*_*ii*_ = −72 pA, *U*_*ee*_ = *U*_*ei*_ = 0.5, *U*_*ie*_ = *U*_*ii*_ = 0.04. Here, the first lowercase index denotes the type (e = excitatory, *i* = inhibitory) of presynaptic neuron and the second index stands for the type of postsynaptic neuron.

Finally, using the above numerical values, let us theoretically determine the regime of synaptic impact and define the concept of a “strong” synapse. To this end, one needs to use the strength-duration curve (SDC) for a single impulse of excitatory synaptic current given by Eq. (14). By its definition, the SDC is a dependence of minimal spike-triggering amplitude 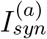 of the stimulating impulse on its characteristic duration, in our case *τ*_*I*_. Assuming that the stimulated non-pacemaker neuron is initially at rest, the SDC is determined by the system of two algebraic equations: *V* = *V*_*th*_ and *dV*/*dt* = 0 at *t* = *t*_*sp*_ [93]. The solutions of this system are 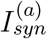 and the moment *t*_*sp*_ of spike generation, as functions of *τ*_*I*_. The SDC is function 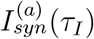.

Let us now consider an idealized case where a non-pacemaker LIF neuron has only one incoming excitatory synapse, which is activated at the moment t = 0, and that the initial conditions are as follows: 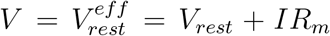 , which is an asymptotic value of the effective resting potential due to background current I *<* I_*c*_ = (*V*_*th*_ − *V*_*rest*_)/*R*_*m*_ (see Eq. (7)), and *x* = *x*_0_, *y* = 0, *z* = 1 − *x*_0_.

Then, according to Eqs. (14) and (15), the synaptic current impulse is given by

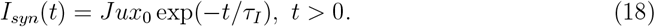

The corresponding SDC for the amplitude 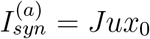 reads [93]

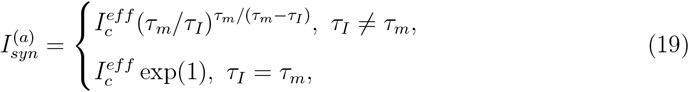

where 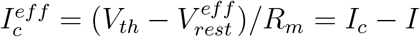. From Eq. (19) we get a threshold value for *J*,

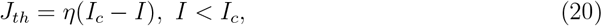

such that for *J* ≥ *J*_*th*_ a synaptic impulse determined by Eq. (18) results in a spike generation by the postsynaptic non-pacemaker neuron. In what follows, we define a “strong” synapse as that having *J* ≥ *J*_*th*_. Importantly, Eq. (20) quantifies the fact that the synaptic strength is not absolute but relative: it depends on the postsynaptic neuron excitability.

In turn, the dimensionless coefficient *η* in Eq. (20) is defined as (for *τ*_*I*_ ≠ *τ*_*m*_)

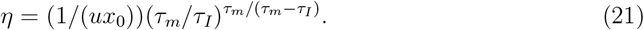

Substituting *τ*_*m*_ = 20 ms, *τ*_*I*_ = 3 ms, and u = *U*_*ee*_ = 0.5 into Eq. (21), one gets *η* ≈ 18.6/*x*_0_. If the background current is zero, *I* = 0, and *x*_0_ = 1, for *I*_*c*_ = 15 pA according to Eq. (20) we get *J*_*th*_ ≈ 18.6I_*c*_ = 279 pA. This value is more than seven times higher than the mean value *J*_*ee*_ = 38 pA (cp. [16]) and exceeds the maximal value 4*J*_*ee*_ = 152 pA allowable in the sampling. So if all non-pacemaker neurons would have zero background currents, then, even without taking into account the synaptic depression, the neuronal network would not contain strong synapses, each of which can independently cause a spike generation by the postsynaptic neuron (see the bottom right graph in Fig. 1). In that case, to make a non-pacemaker neuron fire, several incoming excitatory synapses must be activated in a time interval substantially smaller than the relaxation time constant *τ*_*m*_ of the neuron’s potential (see Eq. (7)). This condition is increasingly unlikely if the fraction of pacemakers is getting smaller.

Nonzero subcritical background currents drastically change the network dynamics. In fact, due to those, the effective (asymptotic) neuronal resting potential 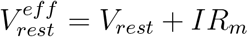 is continuously distributed from *V*_*rest*_ up to *V*_*th*_; this distribution is the same as that for the background currents. It means that a substantial fraction of non-pacemaker neurons can be hypersensitive such that activating even a single incoming excitatory synapse may trigger a spike. This fraction can be quantified as follows.

Provided that J ≤ *J*_ma*x*_ = 4*J*_*ee*_ in Eq. (14) and Eq. (18), from the spike-triggering condition *J* ≥ *J*_*th*_ = *η*(*I*_*c*_ − *I*) one gets the lower value *I*_*_ of the sought range of background currents (the upper value is *I*_*c*_): *J*_ma*x*_ = *η*(*I*_*c*_ − *I*_*_), *I*_*_ = *I*_*c*_ − *J*_ma*x*_/*η*.

Then the fraction of highly excitable non-pacemaker neurons is given by

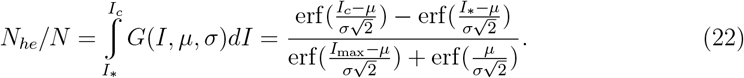

At *x*_0_ = 1, we get *I*_*_ ≈ 6.8 pA and *N*_*he*_/*N* ≈ 0.57, i.e., more than half of all non-pacemaker neurons are potentially highly excitable. But how many of them have at least one strong enough incoming synapse?

Provided that values of *J* in Eq. (14) are normally distributed, the fraction of strong synapses (i.e., those having *J* ≥ *J*_*th*_) for a given *J*_*th*_ ≤ *J*_ma*x*_, or the probability that a given synapse is strong, reads

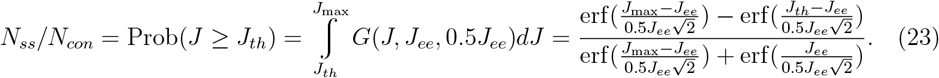

Simplifying the latter formula and substituting *J*_*th*_ from Eq. (20) there, we get

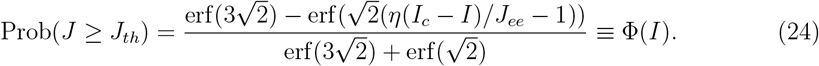

The function Φ(*I*) increases monotonically from Φ(*I*_*_) = 0 to Φ(*I*_*c*_) = 1.

The probability that a highly excitable non-pacemaker neuron has at least one strong enough incoming synapse (note that it is strong enough for that particular neuron) is given by

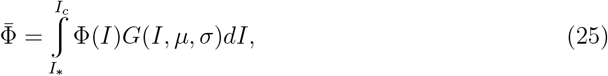

where *G*(*I*, *µ*, *σ*) is standardly determined by Eq. (9). For the above parameters, numerical integration gives 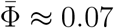.

Accordingly, denoting by *N*_*tr*_ the statistically average number of highly excitable nonpacemaker neurons with at least one strong incoming synapse (in what follows, we call such neurons as “trigger neurons”, see **Discussion**), we get 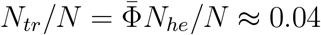. The latter fits well the simulation results described in Sec. 3.4.

It is worth noting a substantial dependence of *N*_*he*_ and 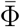 on *x*_0_, i.e., on the initial fraction of synaptic resources in the recovered state. Decreasing *x*_0_ leads to an increase of *I*_*_ in Eqs. (22) and (25) resulting in a decrease in *N*_*he*_ and 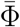 and, consequently, in *N*_*tr*_. For instance, decreasing *x*_0_ by half, from 1 to 0.5, leads to 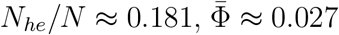 and *N*_*tr*_/*N* ≈ 0.005.

The above theoretical analysis allows us to explain the robustness of the very first population spike that occurs about 30 ms after the simulation begins (see Fig. 2 and Fig. 10, and the description of the initial conditions in Sec. 2.3 below). During this time, the excitability of non-pacemakers increases (accordingly, the number of strong synapses also increases), and pacemakers emit the first spike later than their stationary period 1/*ν* (due to *V*_*reset*_ *≠ V*_*rest*_, or *I*_*r*_ *≠* 0, see Eq. (8)). Apparently, the very first population spike corresponds to the first firing of the majority of pacemakers. Indeed, the corresponding synaptic current pulses from the pacemakers have a maximal amplitude (since *x*_0_ ≈ 1), so the number of strong synapses is also close to the maximum, even though the non-pacemakers have not reached the asymptotic values for their effective resting potentials. Afterwards, for an outgoing excitatory synapse of the pacemaker with self-frequency *ν*, due to synaptic depression *x*_0_ drops from about 1 at the simulation beginning to a stationary amplitude *x*_*a*_ ∼ 1/*ν* (see Eq. (17) and the bottom right graph in Fig. 1). The relative strength of the synapse decreases as well. So the network impact of the first few spikes of a pacemaker is maximal and, consequently, the very first population spike should be considered as an artifact of the model and the initial conditions.

**FIG. 2.**
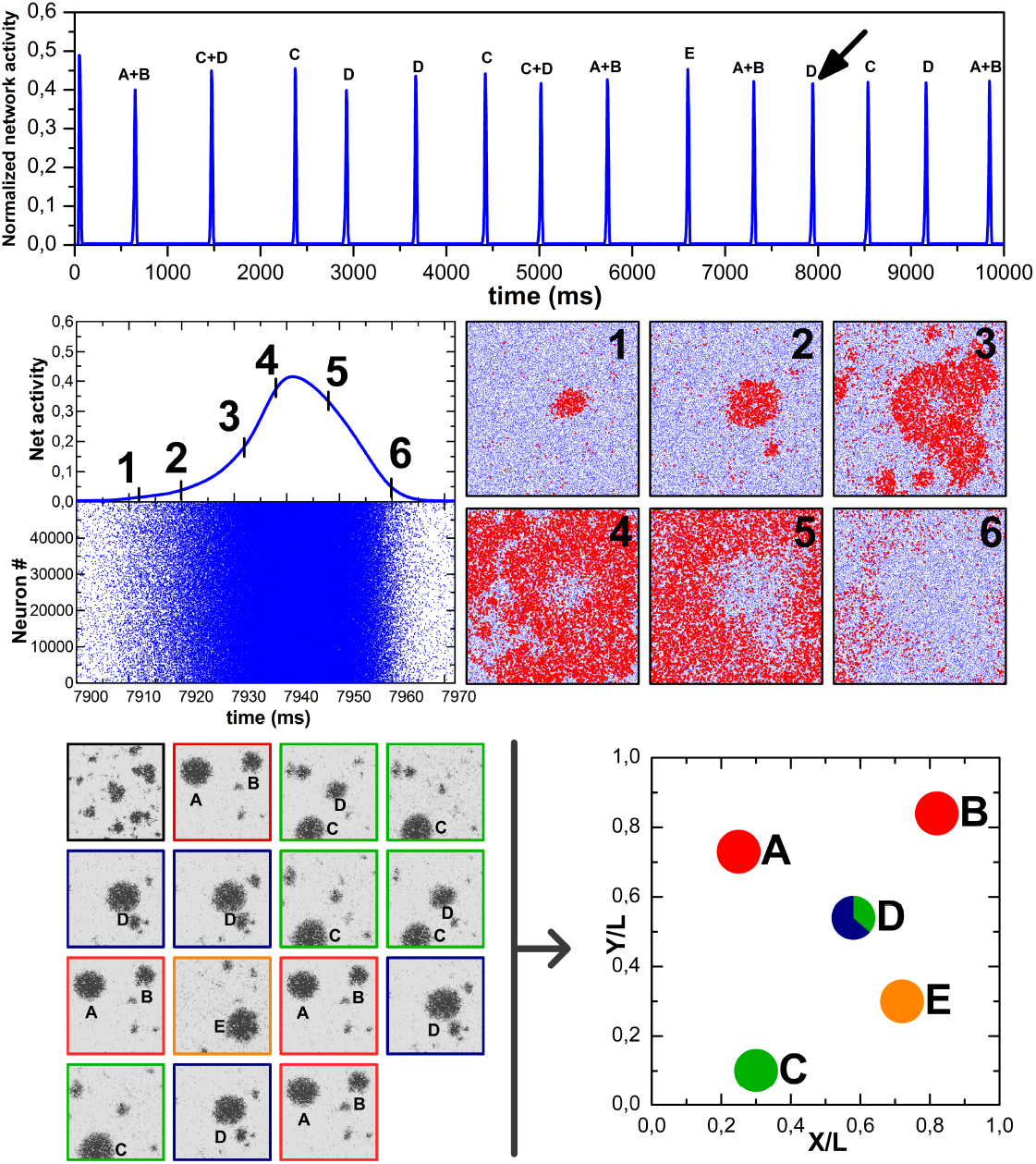
Introducing the reference neuronal network (RefNN) and the spatial map of its n-sites. Upper graph: Time dependence of normalized spiking activity (averaged over 2 ms) of the neuronal network consisting of 50 thousand neurons uniformly distributed over the square area *L × L* at *λ* = 0.01*L*. The peaks of activity are the population spikes. In order to extract a clear spatial map of their n-sites, all inhibitory synapses have been constantly disabled. Capital letters over the population spikes denote specific n-sites generating them (see the bottom right map). Middle graph: LEFT: Network activity (top) and raster (bottom) during the population spike marked by the arrow in the upper graph. RIGHT: Snapshots of the instantaneous spatial activity of neurons for the corresponding moments of the population spike. Blue dots depict neurons and red dots highlight spiking neurons. Each frame corresponds to the whole area *L × L*. On the frame **1** the primary n-site labeled below by D is clearly visible. Bottom graph: LEFT: Spatial locations of the stationary nucleation sites of the population spikes shown in the upper graph. Five primary n-sites (A, B, C, D, E) with different relative rates of population spike generation (see the upper graph) are clearly distinguishable. Black dots depict spatial spiking activity of neurons during 5 ms before and 5 ms after the network activity has crossed the detecting threshold value (*A*_*th*_, see Sec. 2.4). RIGHT: Schematic reconstruction (“map”) of the spatial pattern of n-sites. These are depicted by filled circles, which colors correspond to the different scenarios (e.g., co-activation of two n-sites, C and D) of population spike initiation.

### 2.2. Main output values

The main output values in the numerical simulations are raster, network activity, and spatial coordinates of neurons. The raster plot shows the moments of spike generation for every neuron. In turn, normalized network activity (or, briefly, net activity) A is a histogram showing the number of spikes generated by the network within time bins *△t* = 2 ms and divided by the total number of neurons, *N*. Formally, it is defined as

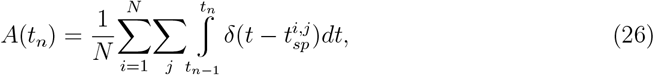

where *t*_*n*_ = *n△*t, *n* = 1, 2, … is the sequence of natural numbers, *t*_0_ = 0, and 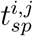 is the moment when *i*-th neuron generates its j-th spike.

Coordinates of neurons and the raster are needed for reconstructing spatiotemporal patterns of spiking activity of the neuronal network (Fig. 2).

### 2.3. Initial conditions and numerical method

The initial conditions for common dynamic variables are the same for all neurons, *V* (t = 0) = *V*_*rest*_, and for all synapses: *x*(*t* = 0) = 0.98, *y*(*t* = 0) = *z*(*t* = 0) = 0.01. For the outgoing synapses of inhibitory neurons, values *u*(*t* = 0) equal to the corresponding U values, which are normally distributed (see Sec. 2.1.3).

All differential equations for the neuronal and synaptic dynamics were solved numerically using the standard Euler method with time step *dt* = 0.1 ms. The numerical simulations have been performed using a custom-made neuronal network simulator written in *C*. The source code of its latest version *NeuroSim-TM-2*.*1* fully compatible with all previous ones is provided (see **Data and code availability**). The code is also available online at https://github.com/dzenn/NeuroSim-TM.

### 2.4. Reconstruction of the spatial pattern of n-sites

The primary n-sites are determined at the initial stage of population spikes. Evaluating the number of n-sites depends on the simulation time, as the population spikes occur randomly from one of them, with different relative probabilities. As a rule, one can obtain the whole stationary set of n-sites after 10-15 sequentially passed population spikes.

A spatial map of n-sites is constructed as follows. The entire area *L × L* is divided into *N*_*elem*_ = 10^4^ elementary square areas with coordinates 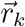. The elementary square side equals 0.01*L*. The spatial map is obtained by counting the number *n*_*k*_ of spikes emitted by the neurons within each of such areas during 35 ms after the moment when the network activity, Eq. (26), is exceeding the specified threshold *A*_*th*_ = 0.006 (for comparison, the baseline activity level typically lies within the range from 0.003 to 0.004). The n-site position is defined as a “center of mass” of 20% of the areas with the largest n_*k*_ values

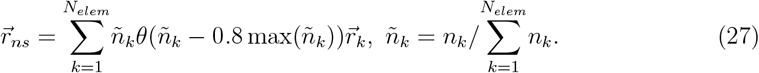

The position of each n-site on the spatial map is shown by a filled circle of the standard diameter 0.12*L*. Each n-site can be assigned a quantitative characteristic the intensity, or relative frequency, of generating population spikes from this site. The intensity of a n-site is depicted by the color saturation of the corresponding filled circle. Thus, for each realization of the neuronal network, a schematic reconstruction of the spatial pattern of stationary n-sites has been made that determines their number, location, and intensities.

Note that values of both threshold *A*_*th*_ and the time window for summing the network activity have been chosen to obtain the most contrast and clear picture of n-sites. However, if a population spike originates from two n-sites simultaneously, the algorithm described above is poorly applicable. Therefore, the automatic determination of all n-sites (using the code *sim data proc*.*c* in the supplementary online files) was additionally controlled visually, by watching videos of spatiotemporal spiking of the network. The videos were generated by custom-made visualization software *Spatial Activity Monitor* [22, 69], rewritten in Python.

## 3. Results

A typical simulation result for the neuronal network model is shown in Fig. 2 (see also Suppl. Video 1). A population spike spatially emerges from one of several stable nsites, from which synchronous spiking activity starts propagating in the form of concentric traveling waves. The spatial map of n-sites is determined by the samplings of simulation parameters, being unique and unchanged for a given neuronal network realization. Each nsite can be quantitatively characterized by its intensity, or relative frequency, of generating population spikes from this site. For a particular network realization, one can therefore build a schematic reconstruction (i.e., a map) of the spatial pattern of n-sites that determines their number, locations, and intensities (Fig. 2, bottom right graph). Extensive simulations have clearly shown that all these characteristics significantly depend on three parameter sets: (1) the sampling of background currents or neuronal excitabilities, (2) the network connectome implementation, i.e., the sampling of the full set of connections between neurons, and (3) the sampling of amplitudes of synaptic currents (*J* in Eq. (14)).

For performing a systematic analysis, we have chosen the network realization, which spiking dynamics is shown in Fig. 2, as a reference neuronal network (RefNN). Each of three numerical experiments described further (Figs. 3–9) is associated with a certain parameter sampling modification of the RefNN and shows the impact of that modification on the pattern of n-sites in comparison to the reference pattern shown in Fig. 2.

**FIG. 3.**
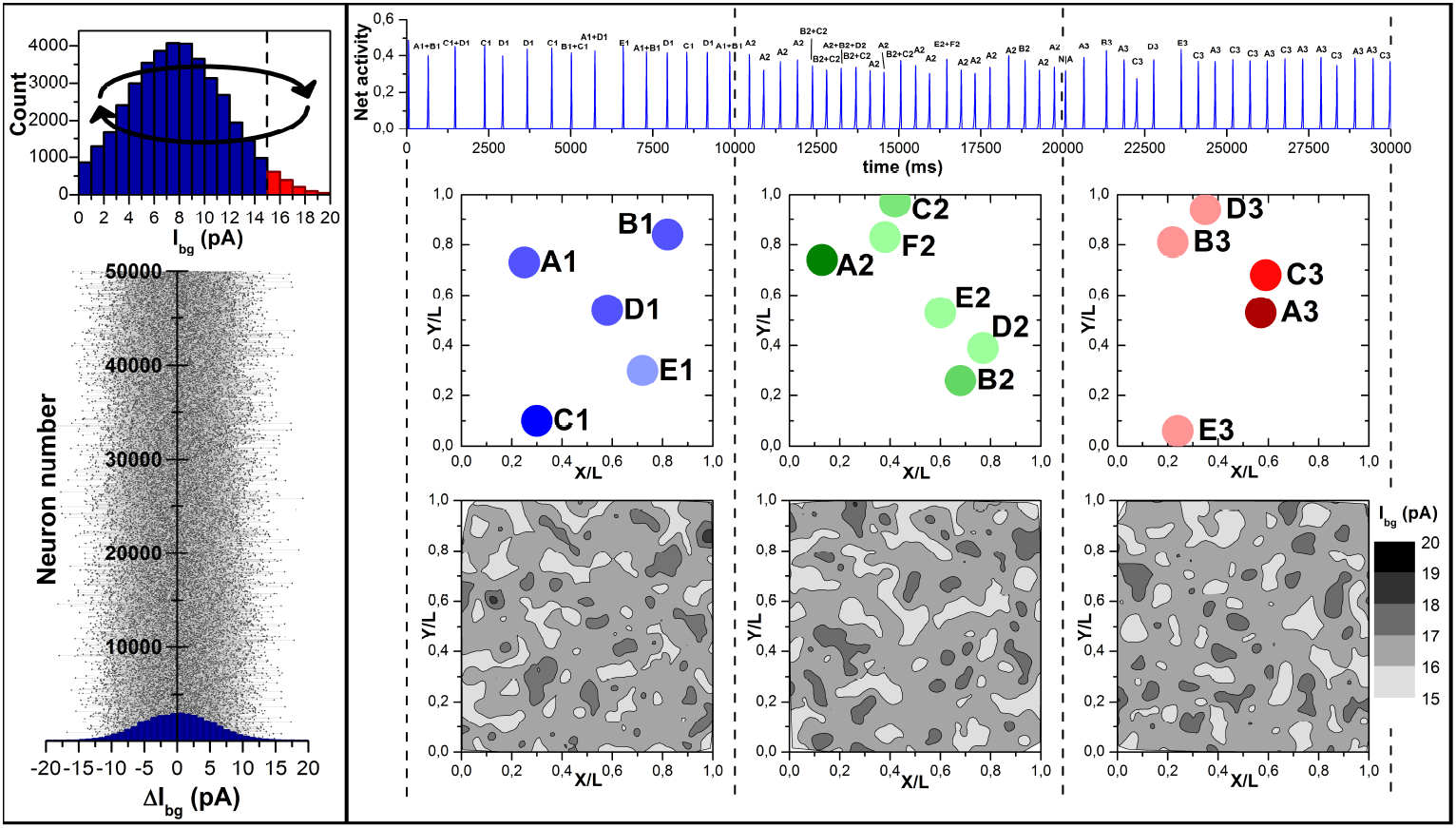
The effect of two random renewals of the entire sampling of background (“bg”) currents for the RefNN, occurring at the 10th and 20th second of the simulation. Left panel: TOP: Network distribution of bg-currents for the RefNN shown in Fig. 2. The bg-currents of pacemaker neurons (i.e. having *I*_*bg*_ *> I*_*c*_ = 15 pA) are marked by red color. The circular arrows show that new values of bg-currents are set independently of the old values so that a pacemaker can become non-pacemaker and vice versa. BOTTOM: A detailed representation of re-sampling the entire set of bg-currents at the 10th second of the simulation. Right panel: The panel is divided into three vertical sections by dashed lines. Each section corresponds to a fixed sampling of bg-currents. TOP: Time dependence of the network spiking activity. Population spikes have been labeled according to their primary n-sites. MIDDLE: Stationary maps of n-sites. The color intensity (saturation) of the circles corresponds to the relative intensity of a particular n-site. BOTTOM: Spatial distribution of pacemakers as a function of the bg-current sampling. During the first 10 seconds both the network activity and the map of n-sites coincide completely with their RefNN benchmark (Fig. 2). The change of the bg-currents sampling, visible by the change of pacemaker density (the lower row of the graphs), leads to a new pattern of stationary n-sites. Note the absence of any correlation between the maxima of pacemaker density and the n-sites.

**FIG. 4.**
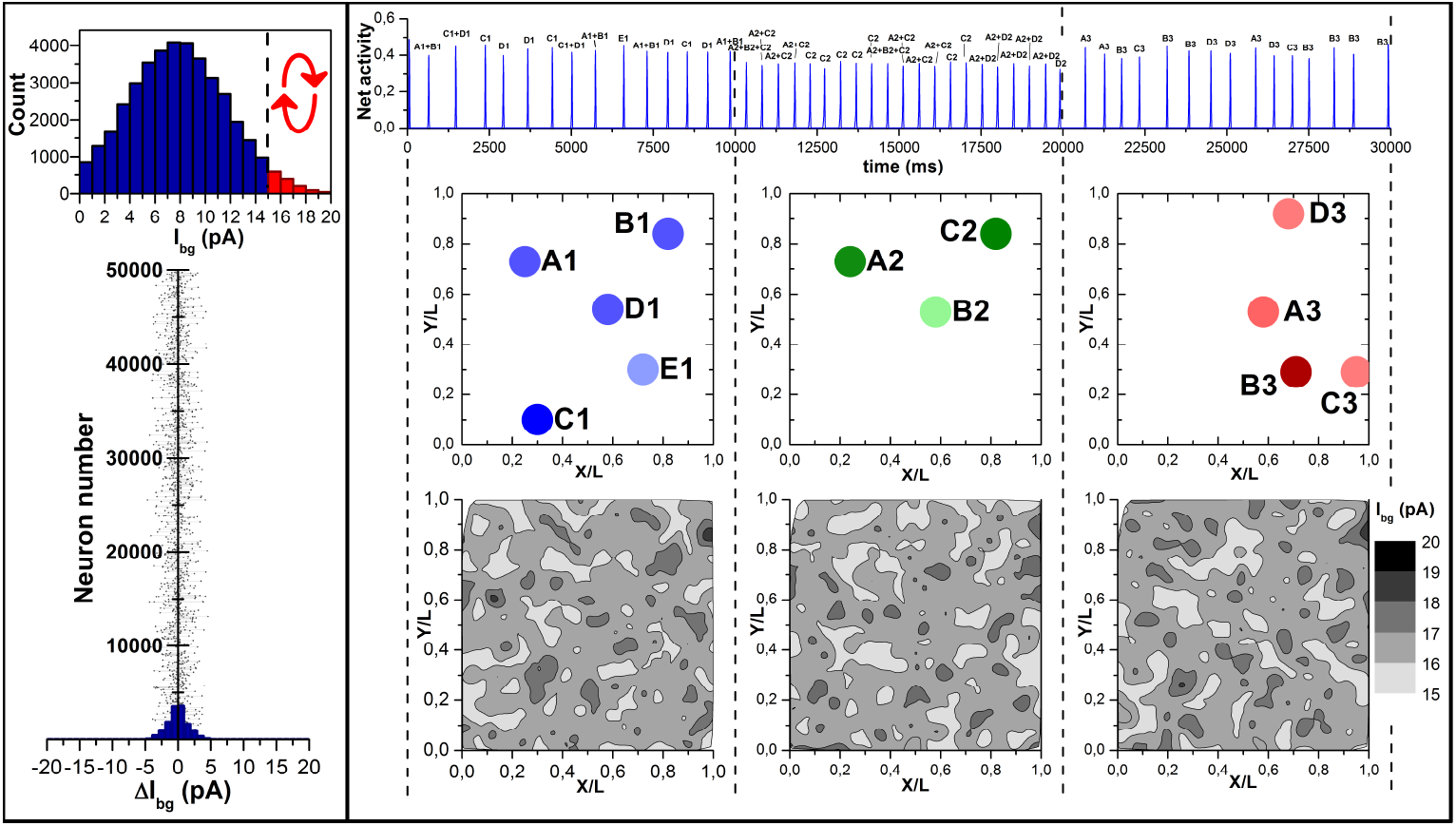
The effect of two random renewals for the distribution of background (“bg”) currents only for pacemakers (3.4 % of the total number of neurons), occurring at the 10th and 20th second of the simulation. Left panel: TOP: Network distribution of bg-currents for the RefNN shown in Fig. 2. Arrows indicate that the redistribution of bg-currents occurs only among pacemakers, so that each pacemaker still remains a pacemaker, but with a different value of the bg-current in the range from 15 to 20 pA. BOTTOM: An expanded representation of re-sampling the bg-currents for pacemakers at the 10th second of the simulation. Right panel: The panel structure is the same as that shown in Fig. 3. In contrast to the case of re-sampling all bg-currents (Fig. 3), the locations of some n-sites (e.g., A1 and A2, B1 and C2, or D1, B2 and A3) were nearly retained.

**FIG. 5.**
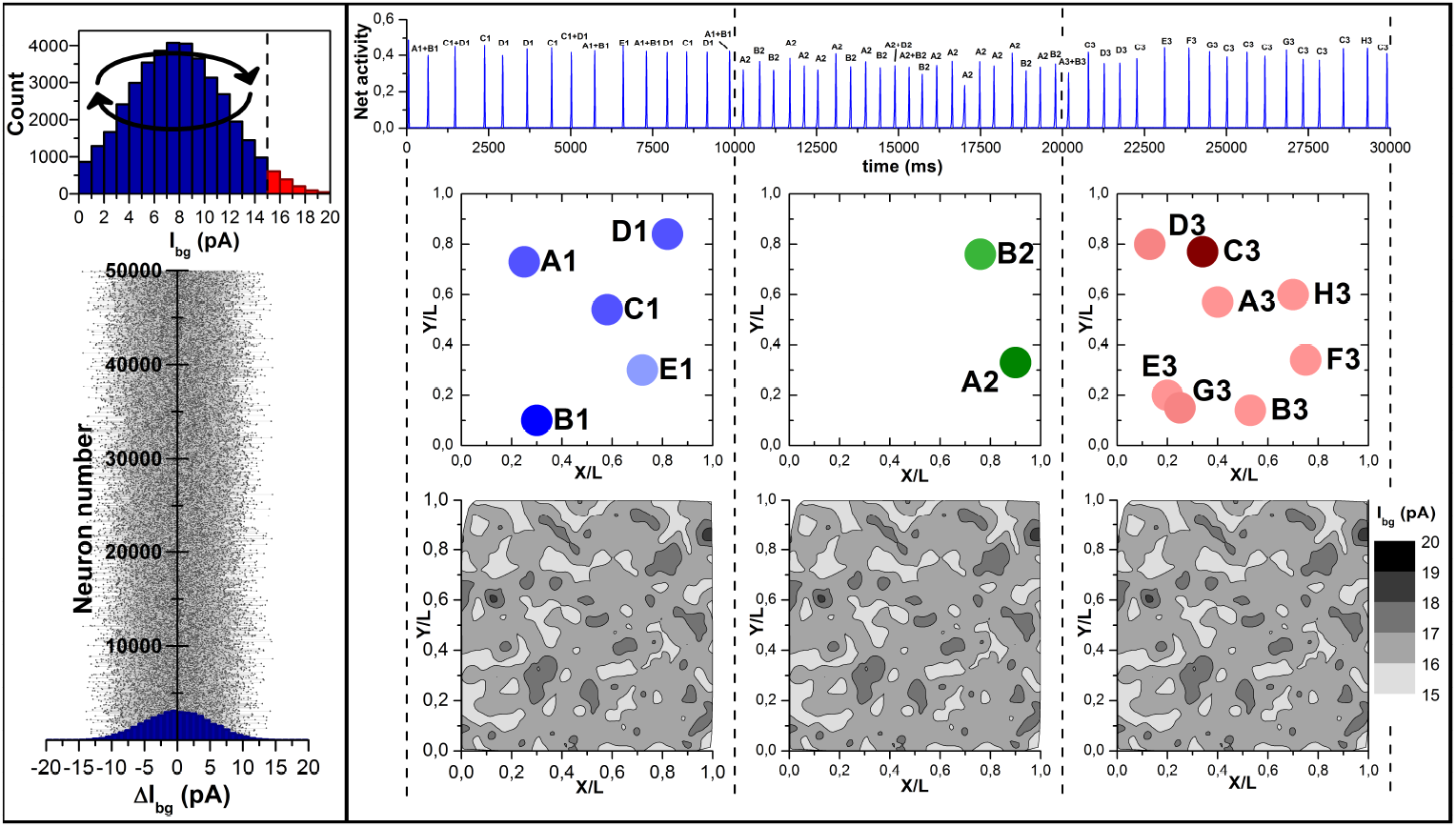
The effect of two random renewals of the sampling of background (“bg”) currents only among non-pacemakers, occurring at the 10th and 20th second of the simulation. Left panel: TOP: Network distribution of bg-currents for the RefNN shown in Fig. 2. Arrows indicate that the redistribution of bg-currents occurs only within the group of non-pacemakers. BOTTOM: An expanded representation of re-sampling the bg-currents for non-pacemakers at the 10th second of the simulation. Right panel: The panel structure is the same as that shown in Fig. 3. Unlike the case of changing the sampling of bg-currents only for pacemakers, the same procedure only for non-peacemakers leads to a substantial change in the number of n-sites. Note that the pacemaker density (the lower row of the graphs) is unchanged.

**FIG. 6.**
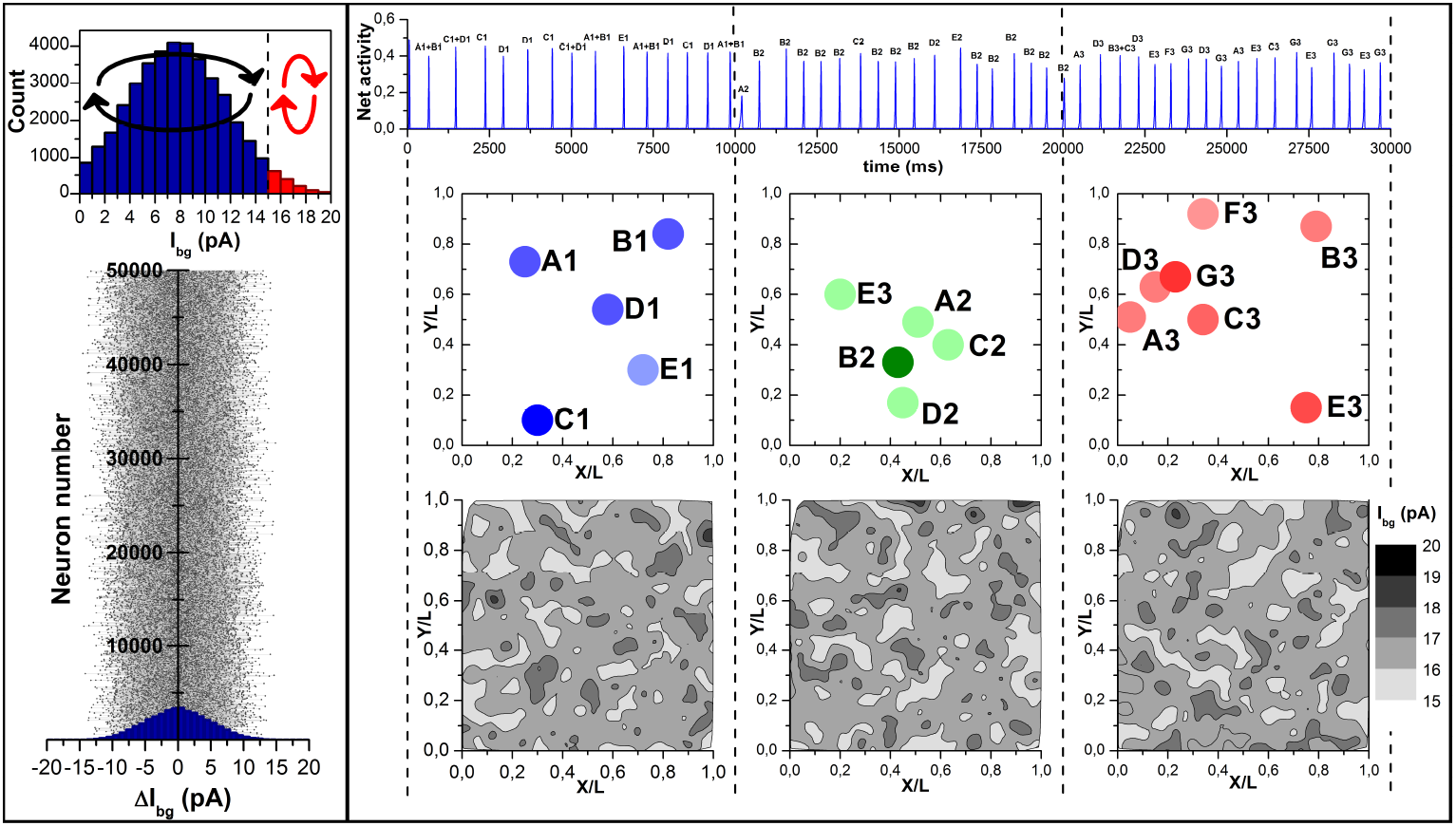
The effect of two random renewals of the sampling of background (“bg”) currents separately among pacemakers and among non-pacemakers (i.e., without mixing between these groups), occurring at the 10th and 20th second of the simulation. Left panel: TOP: Network distribution of bg-currents for the RefNN shown in Fig. 2. Arrows indicate that the redistribution of bg-currents occurs separately within the groups of non-pacemakers and pacemakers. BOTTOM: An expanded representation of re-sampling the bg-currents at the 10th second of the simulation. Right panel: The panel structure is the same as that shown in Fig. 3. It is seen that the pattern of n-sites changes significantly after each re-sampling of the bg-currents.

**FIG. 7.**
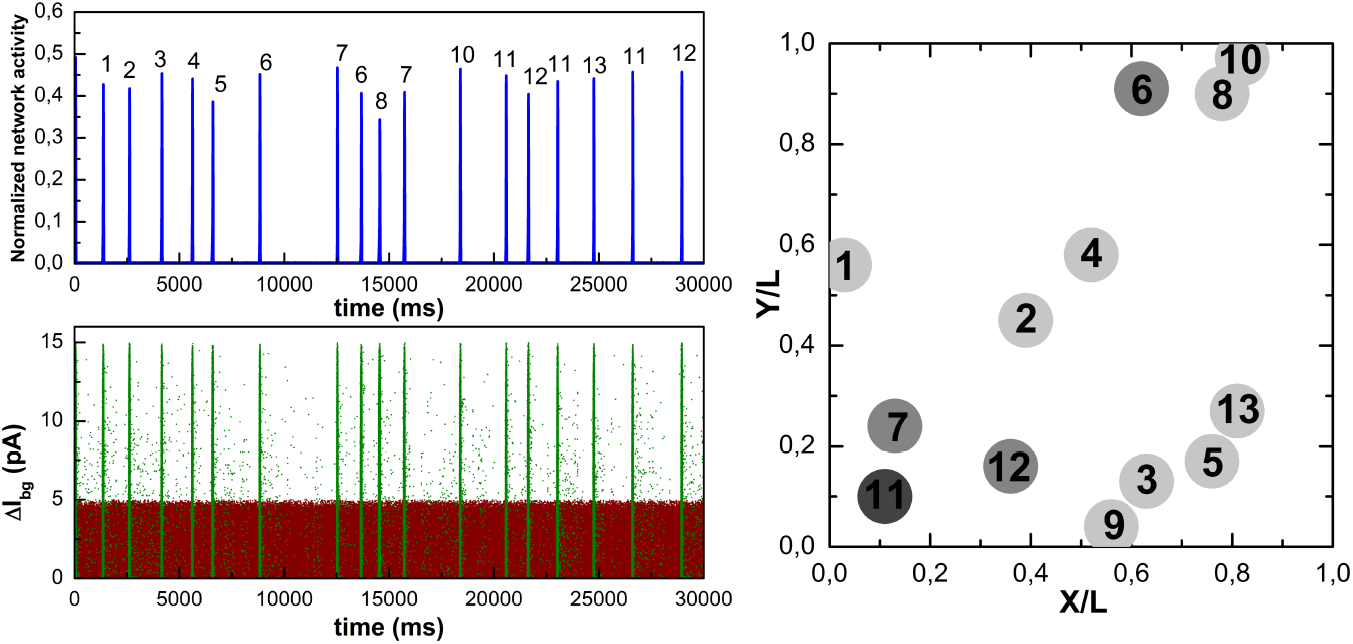
Simulation of spiking activity of the RefNN for the case where the background (“bg”) current value of a neuron is updated each time after generating a spike by this neuron, given that the functional identity of the neuron is retained (i.e., the only possible transitions are “pacemaker → pacemaker” and “non-pacemaker → non-pacemaker”, as for the case shown in Fig. 6). In the present case, population spikes and their nsites still occur, although it happens relatively rarely (note the extended simulation time). Importantly, the location of a currently active n-site is repeated substantially less frequently and only for a few n-sites (see the ones with labels 6, 7, 11, and 12), indicating the absence of a stationary map of n-sites. LEFT: Top graph: Time dependence of the network spiking activity. The number next to each population spike is a label of the corresponding n-site (here, we use numbers as labels because there may be an unpredictably high number of one-shot n-sites). Bottom graph: Absolute values of changes in bg-currents of non-pacemaker neurons (green dots) and pacemakers (dark-red dots) as functions of time. The critical current is *I*_*c*_ = 15 pA and the maximal background current value is *I*_max_ = 20 pA, see Sec. 2.1.2, so the change span for non-pacemakers is from 0 to 15 pA, and for pacemakers it is from 0 to *I*_max_ − *I*_*c*_ = 5 pA. RIGHT: Reconstruction of the spatial locations of the emerged n-sites.

**FIG. 8.**
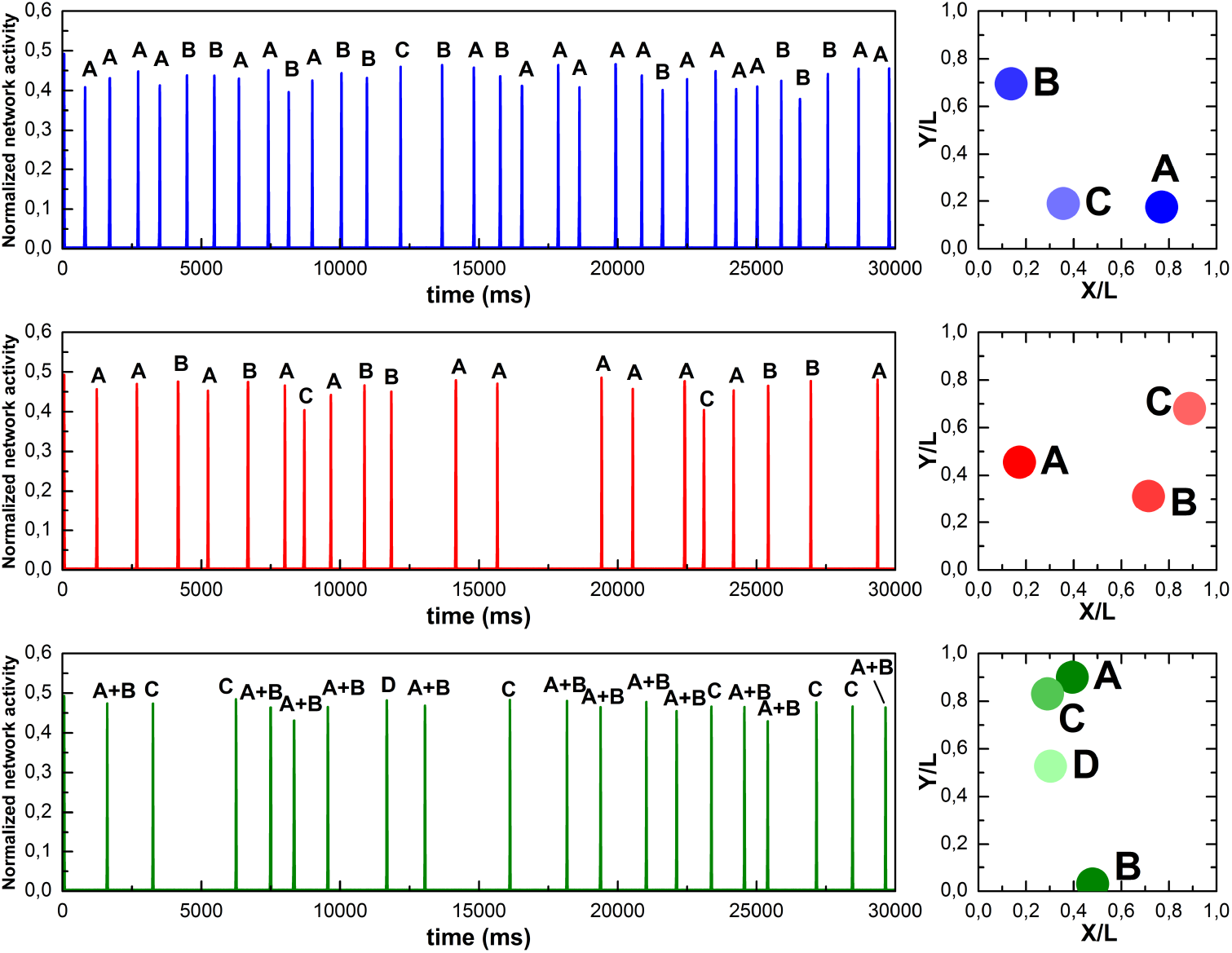
Three simulations (upper, middle and lower panels) of spiking activity of the network, in which (i) the coordinates of neurons and the sampling of background currents are identical to the RefNN, (ii) the synaptic parameters are the same for all synapses of the network and are equal to the mean values of the corresponding Gaussian distributions, and (iii) the network’s connectome is made anew for each simulation. The layout for each panel: LEFT: Time dependence of the network spiking activity. Each population spike is labeled (capital letters A, B, C …) the same as the primary n-site from which it has originated. RIGHT: Reconstruction of the spatial map of stationary n-sites labeled by capital letters. It is seen that the maps are significantly different for the different realizations of the network’s connectome.

**FIG. 9.**
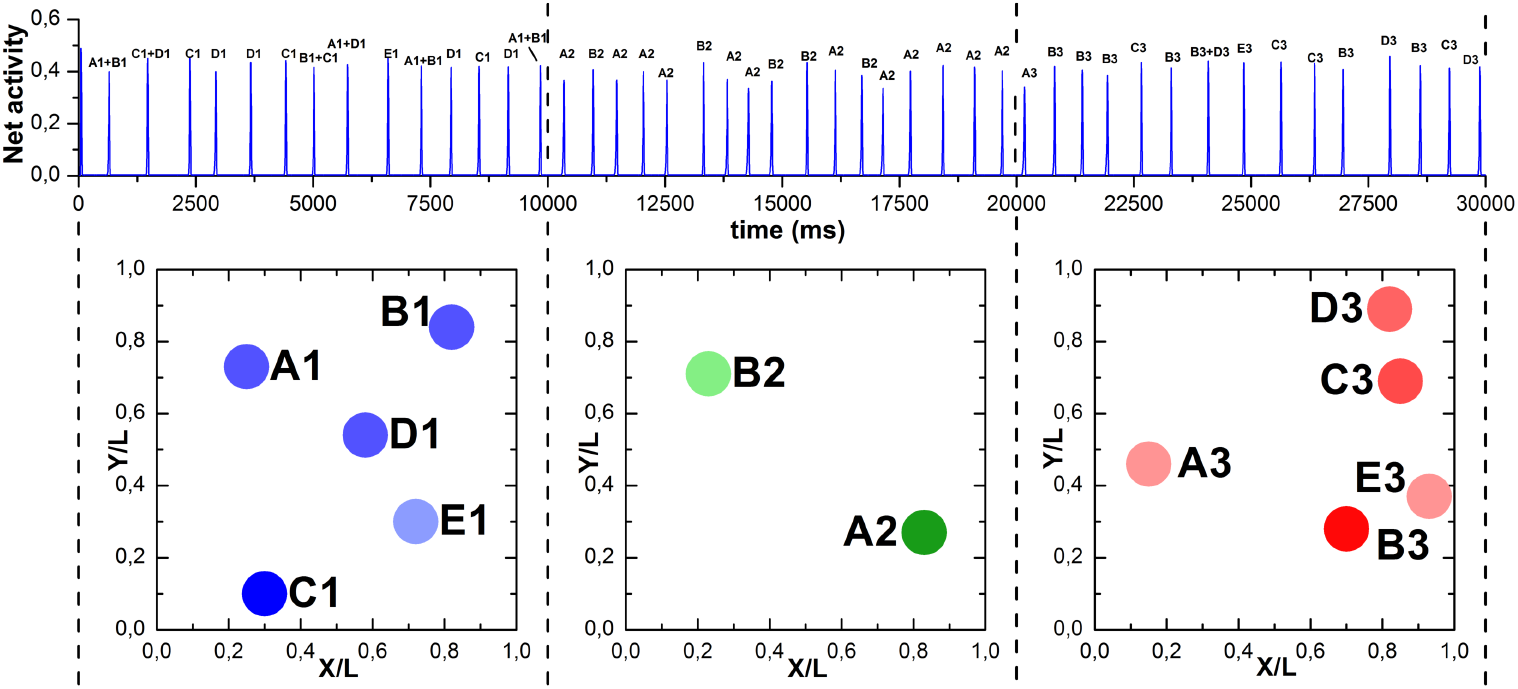
Simulation of the RefNN spiking activity with two instant re-generations (at the 10th and 20th second) of the whole sampling of amplitudes of synaptic currents. Similar to the right panel in Fig. 3, the graph is divided into three vertical sections by dashed lines. Each section corresponds to a fixed sampling of synaptic current amplitudes. Top: Time dependence of the network spiking activity; the population spikes are labeled the same as their n-sites. Bottom: The stationary map of n-sites. As before, the color intensity of the filled circles corresponds to the relative intensity of a particular n-site. It is seen that the map of n-sites changes significantly after each re-sampling of the amplitudes of synaptic currents.

**FIG. 10.**
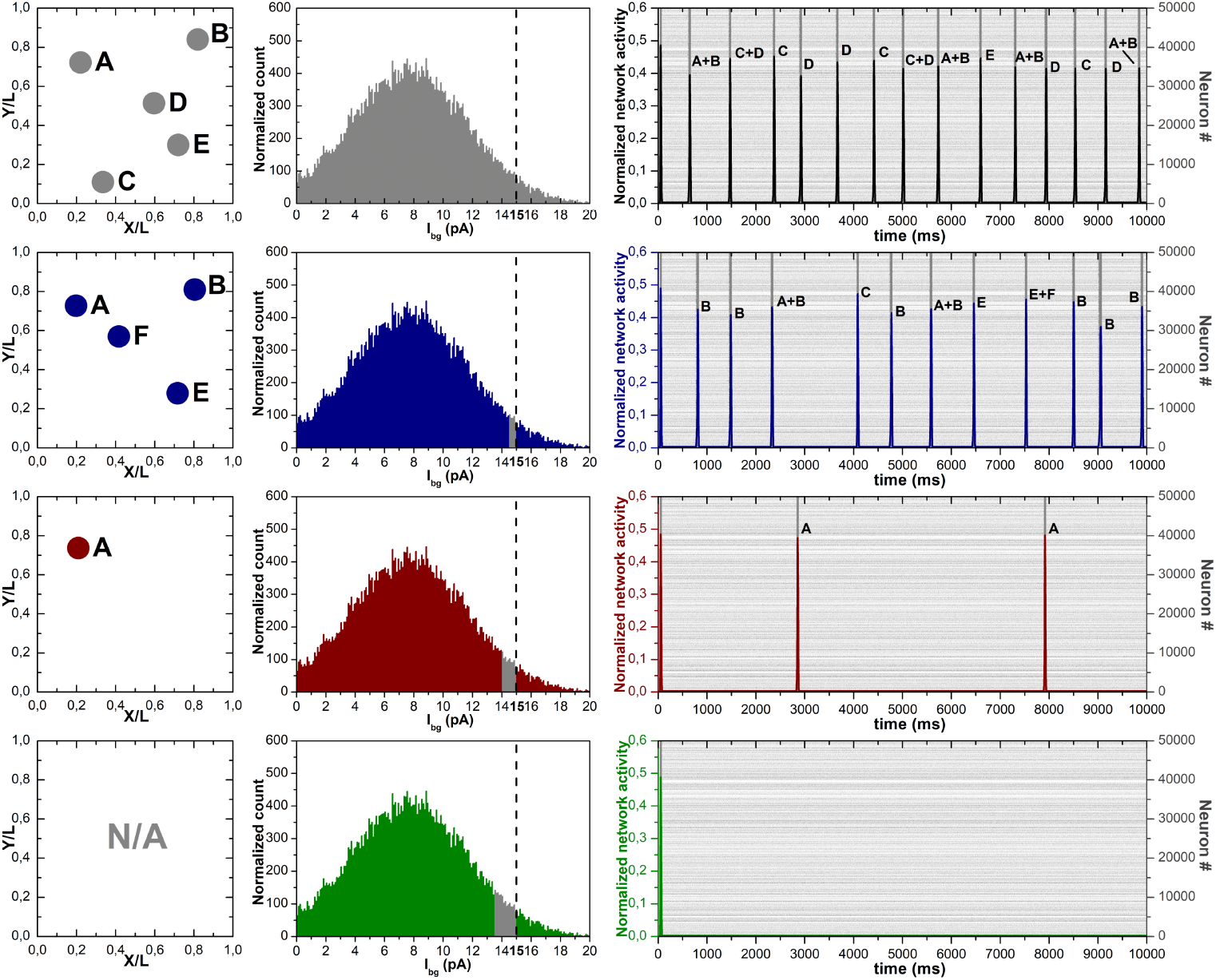
Simulation of the spiking activity for RefNN while blocking the dynamics of non-pacemaker neurons having background (“bg”) current values in a certain range up to the critical value *I*_*c*_ = 15 pA. Top-row graphs (from left to right): the map of n-sites (left), the network distribution of bg-currents (center), and a combined graph of the network activity and the raster plot for the original “unperturbed” RefNN (right), i.e., the same data as shown in Fig. 2. For each population spike, the corresponding “parent” n-site (or a couple of simultaneously active n-sites) is indicated on the activity graph. The 2nd row from the top: similar graphs for the case of blocking the dynamics of neurons having bg-currents ranging from 14.5 pA to 15 pA. This range contains 1.1% of the network’s neurons. Here and further, the blocked range is highlighted by gray color on the distribution of bg-currents (the central graph in the row). Note that the n-sites C and D have disappeared, but a new n-site F has appeared instead. The 3rd row from the top: results for blocking the dynamics of neurons having bg-currents in a wider range from 14 pA to 15 pA, which contains 2.4% of all neurons. In this case, all n-sites, except for site A, have disappeared. Bottom-row graphs: results for blocking the dynamics of neurons having bg-currents in an even wider range from 13.5 pA to 15 pA, which contains 4.1% of all neurons. In this case, all n-sites and population spikes have disappeared (except for the initial-conditions-caused artefact at the beginning of the simulation).

Another important result has been obtained while blocking the dynamics of a small fraction (up to 4%) of neurons with the highest subcritical excitability: it turns out that both n-sites and population spikes are very sensitive to such a blocking, meaning that the most highly excitable non-pacemaker neurons are crucial for the spatially nucleated sync (see Fig. 10).

Finally, we have also thoroughly studied the influence of (i) simulation duration, (ii) heterogeneity of initial neuronal potentials, and (iii) tenfold smaller time step of simulation on completeness and stability of the RefNN n-sites map (see Figs. S2-S4 in Suppl. Mat.).

### 3.1. Changing the sampling of background currents during simulations

The critical current *I*_*c*_ divides the sampling of background currents into two parts such that the minority (a few percent) of neurons with background currents above *I*_*c*_ are pacemakers, and most neurons have background currents below *I*_*c*_ so these are non-pacemakers. It is worth noting that if all outgoing connections from pacemakers to non-pacemakers are blocked, it would lead to a complete cessation of the spiking activity of the latter.

Given that ten seconds of simulation is enough to build the n-sites map, we conducted a series of five experiments on the sampling of background currents, in four of which the sampling was changed in a certain way twice during the simulation, at the 10th and 20th second. In the fifth experiment, the sampling was dynamic all the time: a neuron’s background current was randomly updated after each time the neuron generated a spike. Herewith, in all five experiments the positions of neurons, the network connectome, and the synaptic parameters remained unchanged, i.e., identical to the RefNN realization.

Specifically, in the first experiment, the entire sampling of background currents was generated anew twice, at the 10th and 20th second of the simulation. Such re-sampling could transform a pacemaker into non-pacemaker and vice versa. The experiment result is shown in Fig. 3: the map of n-sites changes significantly with each re-sampling.

In the second experiment, we randomly updated the background currents twice only for the pacemaker neurons: all new values were still above *I*_*c*_ (see Fig. 4). As a result, a few n-sites seemingly remained, with retained or slightly displaced locations. As a result, a few n-sites appeared (or remained) on the former or slightly displaced locations. At the same time, new intense n-sites appeared in other places. Importantly, the locations of n-sites do not correlate with the maxima of the spatial distribution of pacemakers (see the bottom graphs of the right panel in Figures 3 to 6).

In the third experiment, the values of background currents were randomly renewed twice only for non-pacemakers (Fig. 5). Here, unlike the two previous cases, one can clearly see not only the change in locations and intensities of n-sites, but also a significant change in their number.

In the fourth experiment, the background currents were also updated twice, separately within the subgroups of pacemakers and non-pacemakers, so that there were no transitions between the subgroups (Fig. 6). In this case, the map of n-sites was also changed significantly every time and, again, this occurred without any noticeable correlation with the change in the spatial distribution of pacemakers.

Finally, in the fifth experiment, the background current value of a neuron was updated each time immediately after generating a spike by this neuron in such a way that the neuron always retained its group affiliation either to pacemakers or non-pacemakers, as in the fourth experiment. The result is shown in Fig. 7. In this case, population spikes are relatively rare and, most importantly, the locations of n-sites are apparently non-stationary, i.e., population spikes mostly arise from non-repeating n-sites. Abolishing the condition for a neuron to remain within its former group of either pacemakers or non-pacemakers leads to the rapid disappearance of spiking activity in the neuronal network, since the transition from pacemakers to non-pacemakers is much more probable than the inverse one.

### 3.2. Network connectome modification

Despite the specific implementation of the network connectome (i.e., the sampling of the whole set of connections between the neurons) is obviously important for most properties of the network spike activity, we have studied this issue quantitatively. In particular, we performed three simulations based on the RefNN, in which (i) the coordinates of the neurons and the sampling of the background currents were identical to the RefNN, (ii) the synaptic parameters were the same for all synapses of the network and were equal to the mean values of the corresponding Gaussian distributions (see Sec. 2.1.2), and (iii) the network’s connectome was made anew for each simulation. As before, all inhibitory neurons were constantly disabled throughout the simulations. The results are shown in Fig. 8. It is seen that the stationary map of n-sites in every simulation is completely different from the other two. This fact confirms the crucial role of a particular realization of the network connectome for nucleation pattern formation.

### 3.3. Modification of the sampling of synaptic current amplitudes during simulation

In addition to studying the impact of modifying the network connectome, we also conducted a simulation, where, for the RefNN, the sampling of amplitudes of synaptic currents was generated anew twice during the simulation (at the 10th and 20th second). The result is shown in Fig. 9. It is seen that each update of the sampling drastically changes the map of n-sites. It proves that changing only the weights of synaptic connections is sufficient to remake the stationary configuration of the n-sites. Importantly, the pronounced sensitivity of the nucleation map to the sampling of synaptic current amplitudes indicates that a change of that map may be used as a novel visual marker of the impact of spike-timing-dependent synaptic plasticity (on including this in the synapse model, see **Discussion**) on a stationary dynamic state of the neuronal network.

### 3.4. Blocking the dynamics for non-pacemakers having nearly-critical excitability

An interesting result has been obtained while blocking the dynamics of non-pacemaker neurons (by constantly clamping their potential to *V*_*rest*_, regardless of incoming signals) with subcritical values of the background current, specifically, in a narrow range from some value *I*_*cut*_ up to the critical value *I*_*c*_ = 15 pA for turning into a pacemaker. The number of blocked neurons *N*_*cut*_ is explicitly given by formula

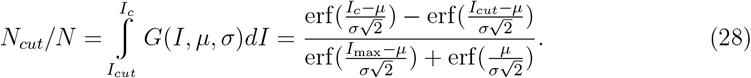

Given that *I*_ma*x*_ = 20 pA, *µ* = 7.7 pA, *σ* = 4.0 pA (see Sec. 2), for *I*_*cut*_ = 14.5 pA, 14 pA, and 13.5 pA one gets *N*_*cut*_/*N* = 1.1%, 2.4%, and 4.1%, respectively. Simulations showed that all n-sites disappeared completely for the latter value, and disappeared partially for two former ones (Fig. 10). In these two cases, the sequence of corresponding population spikes also changed and decreased, relative to the benchmark RefNN simulation (RefNN data are shown on the top-row graphs in Fig. 10). As a result, it turns out that blocking just about 4% of non-pacemaker neurons having the highest subcritical excitability is enough to completely suppress the occurrence of population spikes and their n-sites. A theoretical explanation of this result is given in Sec. 2.1.3.

### 3.5. Testing completeness and stability of the n-sites map: long simulation, initial potentials diversity, and small time step

To test if the simulation duration of 10 sec is sufficient to completely define the stationary map of primary n-sites, we performed a five-times longer simulation for the RefNN. The results are shown in Fig. S2 of the Suppl. Mat. One n-site (labeled by F in Fig. S2) indeed appeared relatively late so it was missed in the simulation shown in Fig. 2. At the same time, most of the n-sites have been revealed within 10 sec of simulation.

We have also checked the stability of the n-sites map against (i) diversity of initial values *V*_*i*_(t = 0), *i* = 1, *N*, of neuronal membrane potentials (see Fig. S3) and (ii) tenfold smaller value of the simulation time step dt defined in Sec. 2.3 (see Fig. S4).

For the case of different initial potentials, we performed five simulations for the RefNN with the duration of 30 sec: the first one (top graph in Fig. S3) was a control simulation with *V*_*i*_(*t* = 0) = *V*_*rest*_, and in each of the four others the values *V*_*i*_(*t* = 0) were randomly taken from the uniform distribution in the interval between the resting potential *V*_*rest*_ and the firing threshold *V*_*th*_. The result is that the diversity of the initial potentials leads to a redistribution of relative activities of previously detected n-sites but not to an emergence of new ones. Notably, the activity alteration is more pronounced for the n-sites, which are relatively faint in the benchmark (“control”) ReNN simulation (Fig. 2).

In turn, to study the influence of simulation time step *dt* on the map of n-sites, we performed three simulations for the RefNN with the duration of 10 sec (Fig. S4). The first one was with *dt* = 0.01 ms (instead of the standard *dt* = 0.1 ms) and the time bin *△t* = 0.2 ms (instead of the standard *△t* = 2 ms) for averaging the network activity defined by Eq. (26). As the result, we got that changing the time step *dt* drastically modified the spatial nucleation pattern: a major part of the benchmark n-sites did not appear, while one new and very active site occurred (Fig. S4). To test if some simple rescaling could revive the benchmark map of RefNN n-sites, in the second simulation, along with the above-mentioned decrease of *dt* and *△*t, we also decreased tenfold the minimal spike-propagation delay *τ*_*del*,*min*_ (see Sec. 2.1.1, at the end), setting it to 0.02 ms instead of the standard 0.2 ms. Finally, in the third simulation, in addition to all changes made in the second simulation, we increased tenfold the spike propagation speed *v*_*sp*_, setting it to 2 mm/ms instead of the standard 0.2 mm/ms. However, all these rescalings did not affect the initial result essentially. We have concluded that a more elaborate rescaling of the integrative properties of neurons is needed to restore the benchmark nucleation map at *dt* = 0.01 ms.

## 4. Discussion

Summarizing, we generated a benchmark implementation of the model neuronal network, called the Reference Neuronal Network (RefNN), and showed a high sensitivity of its n-sites map to

- the sampling of constant background currents during a simulation (Figs. 3-7);
- the network connectome (Fig. 8);
- the sampling of synaptic current amplitudes during a simulation (Fig. 9);
- blocking the dynamics of non-pacemakers having nearly-critical excitability (Fig. 10).

In addition, we checked the n-sites map for the influence of the simulation duration (Fig. S2), the diversity of initial neuronal potentials (Fig. S3), and the small simulation time step (Fig. S4). For the latter, we found a substantial impact on the n-sites map and showed the inefficiency of the two simple re-scalings to offset the impact.

Functionally, the model network exhibits spontaneous population spikes originating from a set of steady n-sites. The crucial question is as follows: what is the structure of a n-site? It is tempting to assume that n-sites are small spatially-localized neuronal assemblies containing a few functionally-equal “trigger” neurons. Firing of one of such neurons, provided that its outgoing synapses are sufficiently strong (see Sec. 2.1.3), may lead to avalanche-like activation of the others in the assembly, i.e., to the n-site activation [17, 94]. That n-site activates other n-sites in the first place and then they collectively activate the rest of the network. A n-site may contain as little as one trigger neuron, though the case where it is formed by several trigger neurons located near each other would make it relatively more stable and active.

In turn, the trigger neurons are thought to be excitatory non-pacemaker neurons having (i) a relatively large number of strong incoming synapses [95] and/or relatively high internal excitability mimicked by the background current value [96] and (ii) a relatively large number of strong outgoing synapses. So being a trigger neuron is a smoothly graded, comparative, but not a binary characteristic. According to the definition, trigger neurons are expected to be systematically active at the start of a population spike generation, and their spiking activity in the intervals between population spikes is expected to be weak, compared to the pacemakers (cp. [8, 97–99]).

It is worth noting that the concept of “trigger” neurons is closely related, or even synonymous, to the neurons of “nacelles” [100], “early-to-fire” neurons [8], “highly active” neurons [101] (cp. [102]), “major burst leaders” [103], “leader” neurons [95–97, 104], “critical” neurons [105], “driver cells” [106], “pioneer” neurons [107], and “core” neurons [108].

Further, our results allow for the interpretation that the stationarity of n-sites can be a quasi-stable (or maybe even unstable) equilibrium position, at least for those n-sites that are not very active relative to the others. It means that even a small change in some of the network model parameters can lead to a substantial reorganization of the whole map of nsites. A potential mechanism for such changes is the Hebbian plasticity, in particular, SpikeTiming–Dependent Plasticity (STDP) [109–112]. The latter can be directly implemented into the model as an additional dynamics of *u*, the fraction of recovered synaptic resource used to transmit the signal across the synapse, in Eqs. (15). Indeed, in the presented model, coefficient *u*, 0 ≤ *u* ≤ 1, has an exact meaning of synaptic weight, being static for the outgoing synapses of excitatory neurons. To include the so-called “online” STDP implementation based on pairwise spike correlations [111], one can add the following dynamic equations for the synapses between excitatory neurons (cp., e.g., [113–115]):

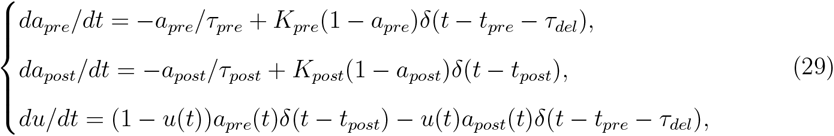

where *K*_*pre*_, *K*_*post*_, *τ*_*pre*_, *τ*_*post*_ are positive constant parameters; *t*_*pre*_, *t*_*post*_ are the moments of the last (up to the current time *t*) presynaptic and postsynaptic spikes, respectively, and *τ*_*del*_ is the spike propagation delay. The auxiliary dynamic variables *a*_*pre*_(*t*) and *a*_*post*_(*t*) are always within the range from zero to one, 0 ≤ *a*_*pre*_(*t*), a_*post*_(*t*) ≤ 1; these trace the spiking history of preand postsynaptic neurons, respectively. Finally, the initial conditions for Eqs. (29) may be as follows: *a*_*pre*_(*t* = 0) = 0, *a*_*post*_(*t* = 0) = 0, *u*(*t* = 0) = 0.5.

For the synapses from excitatory to inhibitory neurons, one could either keep *u*(*t*) = 0.5, i.e., not consider dynamics of synaptic weight u, or allow functionally similar dynamics of *u*(*t*) as for the synapses between excitatory neurons, but with other values of the STDP parameters (cp. [116]). At last, for the outgoing synapses of inhibitory neurons, the dynamics of u is innately described by Eq. (16) in the short-term synaptic plasticity model. One could use parameter U there as a synaptic weight obeyed to STDP. However, as several experiments have shown that inhibitory STDP is strongly dependent on developmental stage of the neural tissue [112], we believe that mathematical modeling of this phenomenon is beyond the scope of this article. Moreover, recent experiments indicate similar developmental-stage dependence even for the “classical” excitatory STDP implied in Eqs. (29) [112]. We therefore emphasize that our consideration of the STDP impact here is purely speculative.

Even though quantifying the impact of the “classical” excitatory STDP on the spatial map of n-sites is a complex unsolved problem [117], the obtained results (see Fig. 9) allow for predicting a substantial influence, provided that STDP substantially affects synaptic weights. As a whole, in the presence of STDP, the n-sites map might require a long transient time to become stationary. For the STDP model described by Eqs. (29), if parameters *K*_*pre*_ and *K*_*post*_ are small enough, the stationary n-sites are assumed to correspond to a stationary bell-shaped distribution of synaptic weights, as this distribution is typical for the STDP rule with multiplicative weight dependence [118, 119]. STDP could also result in a finite lifetime for each n-site, which would blossom and then fade out, with or without further revival.

Another mechanism that may affect the existence and stability of the n-sites map is spike-frequency (or spike-triggered) adaptation of the neuronal dynamics [120–126] (see also a recent model [127] of combined metabolically-driven pacemaking and adaptation). Unlike STDP, this mechanism is internal, i.e., it does not directly depend on interactions between neurons. In the simplest case, the adaptation can be implemented as follows: instead of Eq. (7) for neuronal potential *V* , we now have

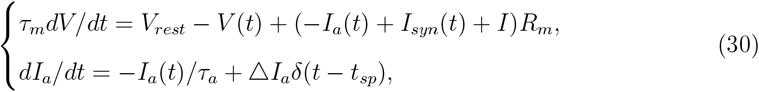

where the second equation describes the dynamics of an adaptation current *I*_*a*_ providing a negative (inhibitory) feedback after each spike of the neuron. Here *τ*_*a*_ and *t*_*sp*_ are the adaptation time constant and the moment of the last generated spike, respectively. Qualitatively, at the moment of spike generation by the neuron, *I*_*a*_ is instantly increased by the value of *△I*_*a*_ and then decays exponentially with characteristic time *τ*_*a*_, which is substantially larger than the absolute refractory period, *τ*_*ref*_ . If incoming synaptic current *I*_*syn*_(*t*) is set to zero and the constant background current *I>I*_*c*_, i.e., the neuron is an unperturbed pacemaker, adaptation current *I*_*a*_ results to a decrease in the firing frequency, compared to that determined by Eq. (8). Importantly, the spike-frequency adaptation and synaptic depression may jointly lead to a nontrivial form of amplification of the impact of pacemakers on other neurons: a decrease in the spike generation frequency due to the adaptation leads to a decrease in synaptic depression (see Eq. (17) and subsequent text) which, in its turn, causes an increase in the amplitude of outgoing synaptic current pulses.

As our simulations have shown a high sensitivity of the n-sites map to pacemaker frequencies (see Figs. 3, 4, and 6), adding *I*_*a*_ dynamics in the spiking network model virtually guarantees some change in the n-sites map relative to the one for RefNN. Moreover, as the adaptation timescale *τ*_*a*_ ∼ 100 ms (cp. [108, 128]) has the same order of magnitude as a typical duration of population spike (see Fig. 2), the adaptation may also directly affect the collective dynamics of non-pacemaker neurons.

Although the effect of adaptation should be studied separately, we assume that it is not similar to the renewal of the background current value each time immediately after spike generation by the neuron (see Fig. 7 and the related text), and the adaptation does not destroy a stationary map of n-sites. Indeed, in the case of the background current renewal immediately after the spike, the new current value can be greater than the old one (except for the two limiting values), while in the case of adaptation the sum (−*I*_*a*_(t) +*I*) in Eq. (30) is always smaller than *I*. In addition, the intervals between population spikes are about ten times longer than the population spike duration and *τ*_*a*_. Therefore, the adaptation currents of non-pacemakers decay significantly during such an interval, and the excitabilities of these neurons will again be determined by the same values of background currents. The n-sites appear at the initial stage of a population spike when most non-pacemaker neurons have almost zero adaptation currents. If the trigger neurons participate in the appearance of nsites only once, i.e. only the first spike from each of them is critical, the effect of adaptation is unlikely to be very significant.

If inhibitory neurons are not blocked in the RefNN simulation, their activity blurs the n-sites map so that it is hardly possible to get a clear map not only algorithmically but even visually (see Suppl. Video 2). In this case, only one or two most intense n-sites corresponding to the map obtained in the inhibition-off mode could be still detectable for the unaltered simulation time. As a rule, inhibition tears local n-sites into several delocalized ragged patches of activity. One can interpret it as follows. In the network model that underlies simulations, the inhibitory neurons have the same spatially uniform distribution and, importantly, the same connectivity characteristics as the excitatory neurons. In real networks, however, inhibitory neurons statistically have a greater proportion of short connections than excitatory ones (e.g., see [59]). This may explain the above-mentioned delocalizing effect caused by the activity of inhibitory neurons. Interestingly, as it has been previously shown in simulations for the same model, in the inhibition-on mode a population spike can occasionally originate in the form of a multi-armed spiral wave with the drifting center if the network connectome is relatively dense (λ = 0.04*L*) [22].

It should be noted that both the neuronal network model and the above-described results on n-sites can be readily extended into the three-dimensional (3D) case, where population spikes have been recently observed in spherical cerebral organoids obtained from human induced pluripotent stem cells [129–131]. Although cerebral organoids have a substantially more deterministic connectome than a model network of randomly connected neurons [132, 133], the latter could be a starting point, as a very rough approximation. Besides, moving from a two-dimensional plane to a three-dimensional space always leads to new possibilities. Indeed, is it possible to make four equilateral triangles out of six matches? On the plane no, but in the three-dimensional space yes (imagine a tetrahedron, i.e., triangular pyramid).

Formally, for the extension into the 3D case, one should only change accordingly (i) the dimensionality of the metric distance and (ii) the probability density *P*(*r*) of finding two neurons at 3D distance *r* from each other. Below, we briefly outline the most basic cases of the cube and the ball, compared to their 2D analogs.

For neurons being uniformly distributed in the cube *L × L × L*, the probability density *P*_*cube*_(*r*) of finding two neurons at 3D distance *r* from each other is given by (as in Eq. (3) before, we equate *L* = 1 to shorten the formula)

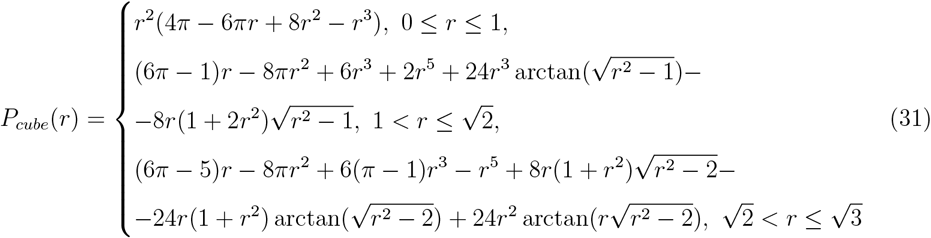

with 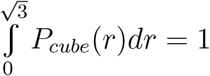 [134] (cp. [135]). The dependence *P*_*cube*_(*r*) is shown by the blue curve on the left graph in Fig. 11.

**FIG. 11.**
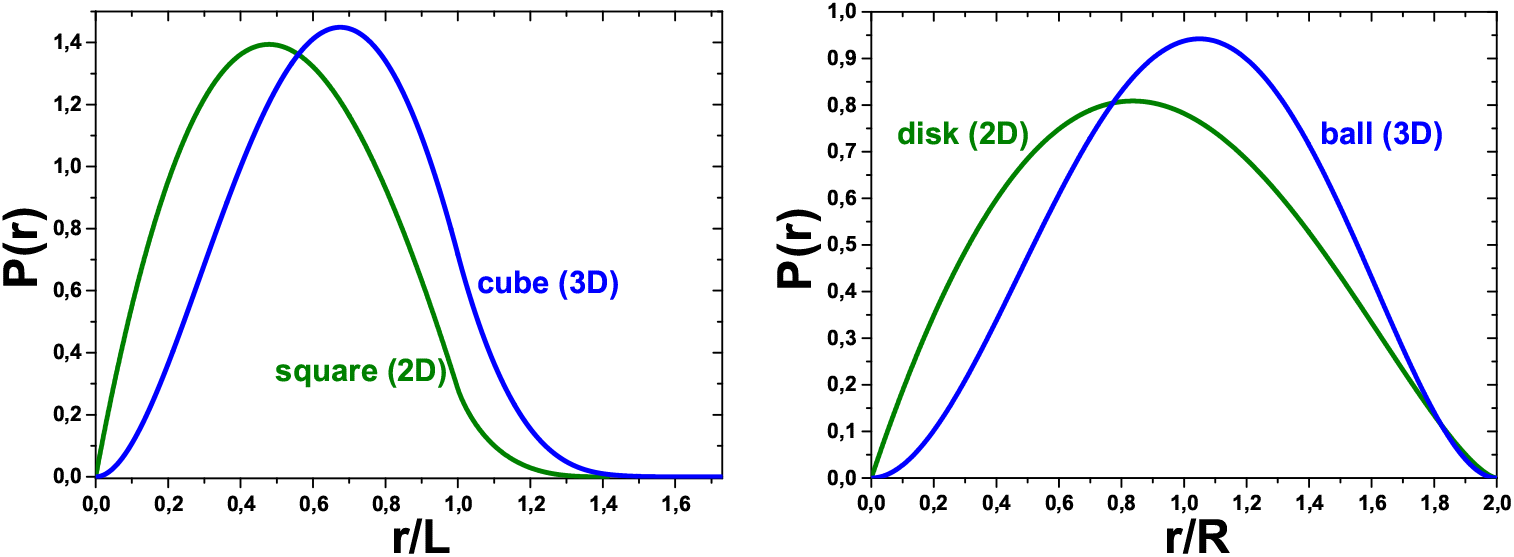
Probability density *P* (*r*) of detecting two points (in our case, two point neurons), randomly and independently dropped in a specific enclosed area (green curves) or volume (blue curves), at the distance *r* from each other. The distance *r* is two-dimensional (2D) for the area and three-dimensional (3D) for the volume. Left graph: The green curve is *P* (*r*) for the square *L × L* (see Eq. (3)), and the blue curve is *P* (*r*) for the cube *L × L × L* (Eq. (31)), where *L* is the corresponding side length. Right graph: The green curve is *P* (*r*) for the disk *πR*^2^ (Eq. (32)), and the blue curve is *P* (*r*) for the ball (4*/*3)*πR*^3^ (Eq. (33)), where *R* is the corresponding radius.

For a centrosymmetric spatial boundary (disk, ball), similar formulas for the probability density *P*(*r*) are much simpler. In particular, for the point neurons being uniformly distributed within a 2D disk [21] and a 3D ball of radius R, the corresponding probability densities *P*_*disk*_(*r*) and *P*_*ball*_(*r*) are given by (see the right graph in Fig. 11)

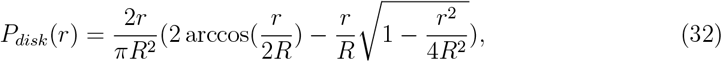

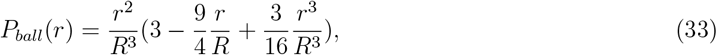

where 0 ≤ *r* ≤ 2*R* and 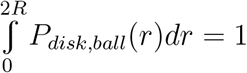 [136–138].

For the 3D case, though a detailed statistical study similar to the one reported here for 2D networks has not been carried out so far, a series of test simulations for 3D neuronal networks confined in the cube has confirmed the existence of population spikes occurring from a few static n-sites.

Based on all of the above, as the spiking network model described here has demonstrated a very high richness-to-complexity ratio, it may be used as an excellent illustrative tool for teaching network-level computational neuroscience, complementing a few benchmark models (e.g., the Brunel’s model and its amended spatial versions [139, 140]). Specifically, to illustrate spatiotemporal patterns of network spiking activity, such as concentric traveling waves underlying population spikes.

Notably, population spikes have been often treated as neuronal avalanches, the statistics of which exhibits signatures (e.g., a specific power-law distribution of the avalanche sizes, cp. [141]) of self-organized criticality (SOC) [142–144]. Moreover, modeling studies have indicated that short-term synaptic plasticity may play a leading role in this phenomenon [145, 146] (reviewed in [147]). In turn, our findings show that population spikes can have a small set of steady n-sites rather than occurring spatially at random. The latter, and that some n-sites can be located quite close to each other, challenge the standard SOC interpretation, which implies a certain spatial symmetry. Further, our results are consistent with those claiming the existence of a small fraction of trigger neurons, rather than a transient formation of neuronal assemblies. Such neurons are crucial for driving population spikes, requiring further elaboration of the SOC-based explanation.

## 5. Conclusion

In this article, for a two-dimensional generative network of excitatory spiking neurons with short-term synaptic depression, we have numerically studied the sensitivity of the spatial map of spontaneously formed n-sites of population spikes to the changes in the network distributions of (i) neuronal excitability, (ii) the set of incoming and outgoing connections of a neuron, and (iii) amplitudes of pulsed synaptic interaction between neurons. We have also detected a crucial role of a few percent of non-pacemaker neurons with the highest excitability both for the nucleation effect and population spikes.

A possible influence of the Hebbian plasticity (STDP) and spike-frequency adaptation on the existence and stability of the n-sites map, and an extension of the network model into the three-dimensional case, have been discussed. An amplification of the impact of pacemakers on other neurons due to an interaction between the spike-frequency adaptation and synaptic depression is qualitatively predicted. Finally, using the example of a simulation with unblocked inhibitory neurons, their blurring impact on the n-sites map has been demonstrated.

In turn, the impact of the outlined results may be as follows. From a theoretical perspective, the proven high sensitivity of the n-sites map to the changes in parameter samplings makes it impossible to describe the nucleation effect using a standard mean-field approach. Interpreting the nucleation as a critical dynamic state is competed by the hypothesis of a decisive impact of a small fraction of trigger neurons having relatively high (i) internal excitability and/or number of strong incoming excitatory synapses and (ii) number of strong outgoing excitatory synapses.

From a practical perspective, our results indicate that instead of adjusting the closedloop control system of a cyborg to the emergent pattern of n-sites, one should find a way to completely suppress its occurrence by minimal, extremely selective exposure (e.g., physical – by laser ablation, optogenetic or, perhaps, pharmacological) to the disinhibited neuronal culture. The targets for such an exposure may be up to 5% of the most excitable nonpacemaker neurons.

Finally, the obtained results, including the complete dataset of simulations, also provide the empirical basis for constructing a theory that would predict the number and locations of primary n-sites without performing simulations of neuronal dynamics, i.e., only based on the static parameter samplings of a particular implementation of the neuronal network.

## Acknowledgements

A.P. is grateful to Chaitanya Chintaluri, Douglas Feitosa Tomé, and Tim Vogels for useful discussions.

## Data and code availability

The attached Supplementary Material for this article contains four Figures, S1-S4, and two Videos. The simulation data and program codes required for reproducing all results of this study are available online at https://doi.org/10.6084/m9.figshare.24527848.

## Author contributions

D.Z. designed and made most program codes, performed simulations, analyzed data, prepared most simulation-based graphs and videos, and verified the manuscript. A.P. conceived the study, designed graph sketches, analyzed data, prepared model-illustrating graphs, performed a theoretical analysis, and wrote the manuscript.

