## Supplementary Figure S1 for "The vitals for steady nucleation maps of spontaneous spiking coherence in autonomous two-dimensional neuronal networks"

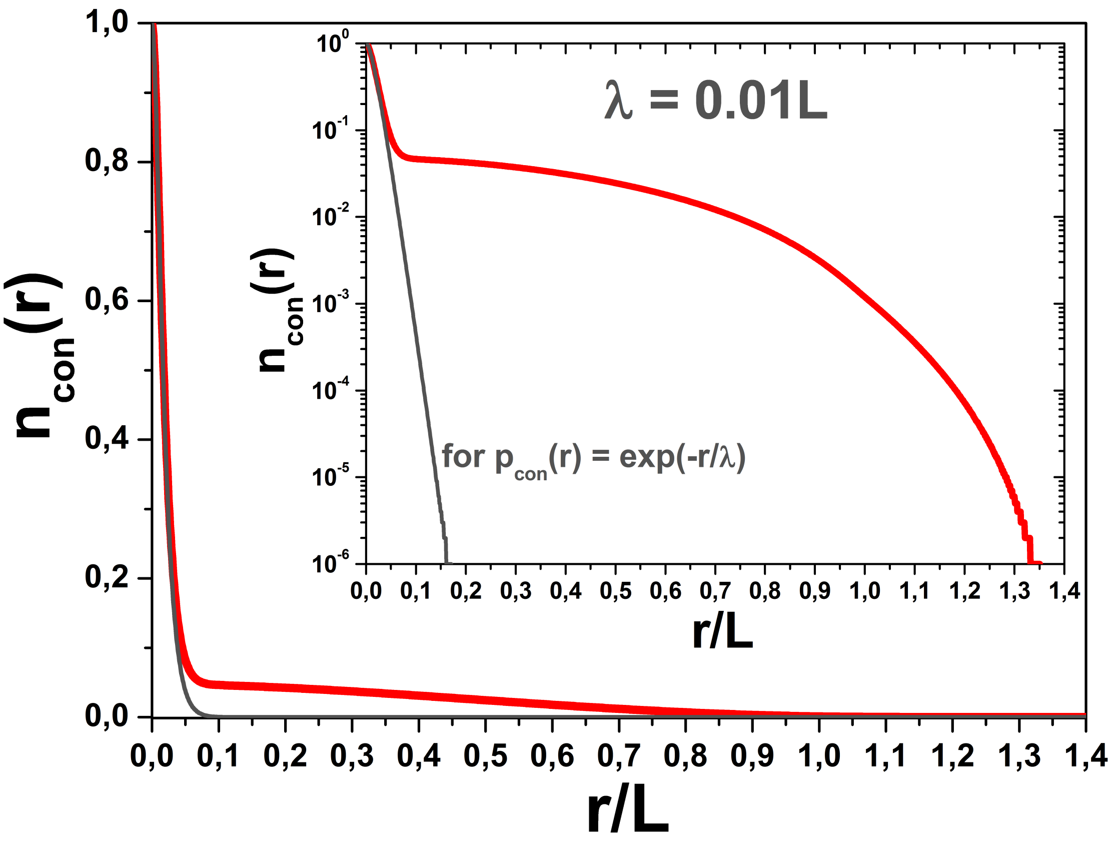


**Figure S1**. The red curve is the fraction $n_{con}(r)$ of network connections with a length longer than $r$; see the formula below or Eq. (5) in the main text. The neurons are uniformly distributed over the square $L \times L$ with probability density $P(r)$ to find two neurons at distance $r$ from each other, see Eq. (3) in the main text. The connection probability is $p_{con}(r) = \exp(-r/\lambda) + p_{\min}\theta(r - r_0)$ with $r_0 = \lambda \ln(1/p_{\min})$. For $p_{\min} \approx 3 \cdot 10^{-5}$ and $\lambda = 0.01L$ used in all simulations, one gets $r_0 \approx 0.1L$.

$$n_{con}(r) = \int_r^{\sqrt{2}} p_{con}(r')P(r')dr' / \int_0^{\sqrt{2}} p_{con}(r')P(r')dr'.$$

The gray curve illustrates the case $p_{con}(r) = \exp(-r/\lambda)$ resulting from $p_{\min} = 0$.

The inset shows the same graph with a logarithmic scale on the vertical axis.
