## Supplementary Figure S2 for "The vitals for steady nucleation maps of spontaneous spiking coherence in autonomous two-dimensional neuronal networks"

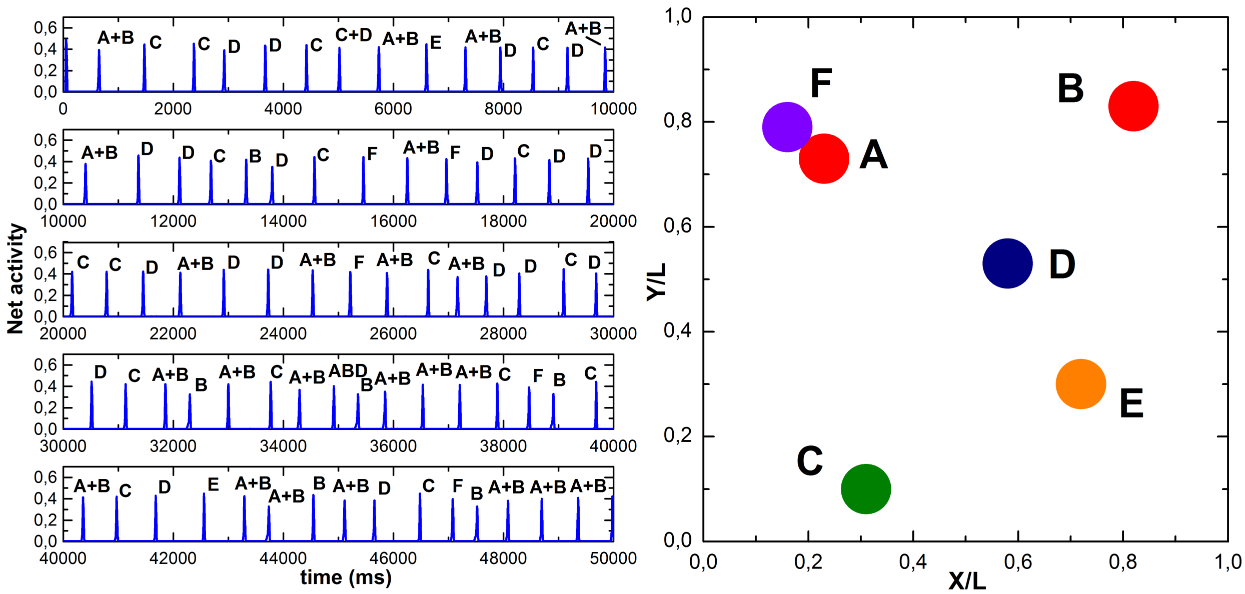

**Figure S2**. Simulation of spiking activity of the RefNN (Reference Neuronal Network) for a long time (50 sec vs standard 10 sec; see Fig. 2 in the main text) to trace the occurrence of any new primary n-sites. As before, the inhibitory neurons (20%) have been blocked all the time.

LEFT: Network spiking activity chunked into five vertically stacked graphs for 10 sec each. As in Fig. 2, the capital letters over the population spikes denote specific n-sites generating them.

RIGHT: Schematic reconstruction of the spatial pattern of primary n-sites. It can be seen that a new n-site F in the vicinity of n-site A does indeed appear relatively late (after 15 sec) so it is not visible in the simulation shown in Fig. 2, which lasted only 10 sec.
