## Supplementary Figure S3 for "The vitals for steady nucleation maps of spontaneous spiking coherence in autonomous two-dimensional neuronal networks"

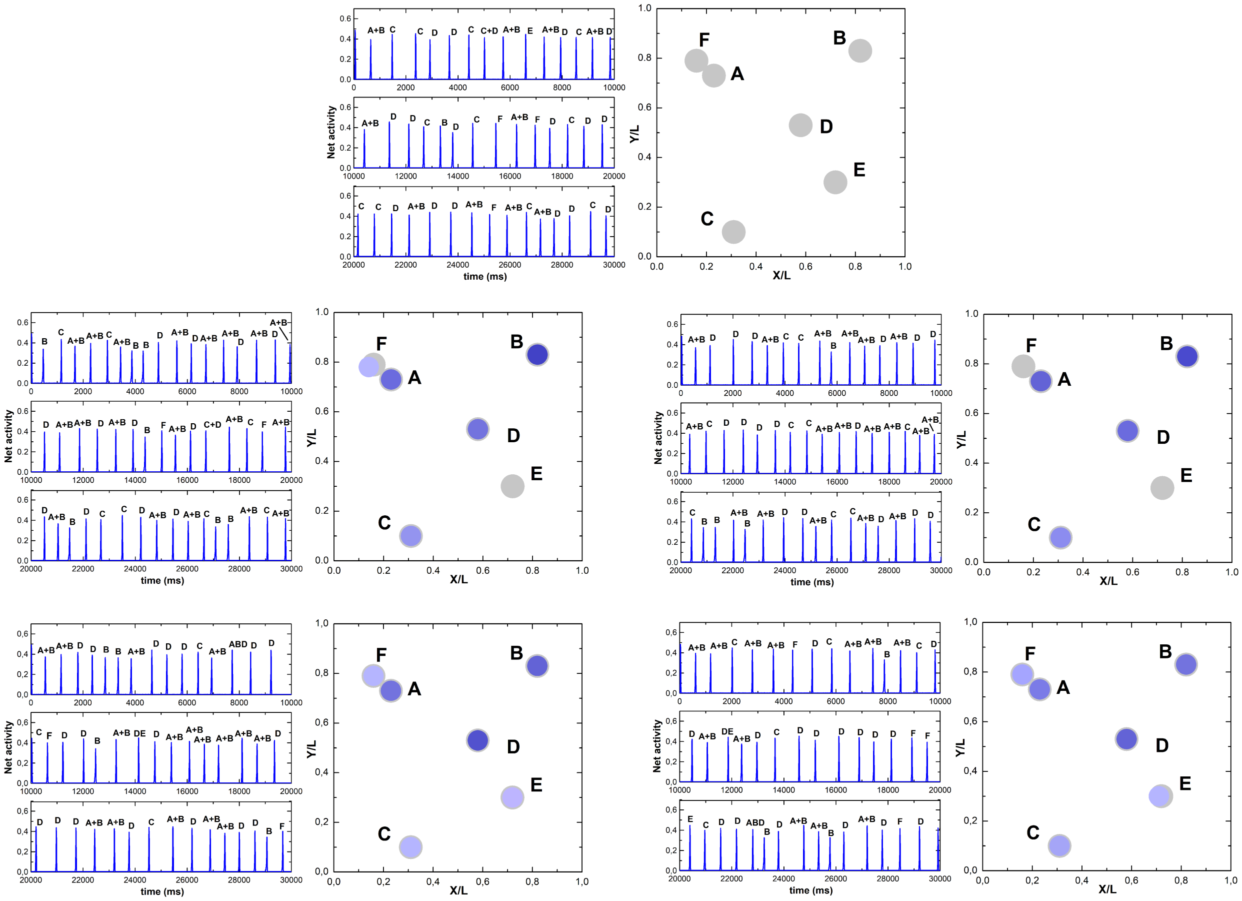


**Figure S3**. Influence of diversity of the initial values $V_i(t = 0)$, $i = \overline{1, N}$, of neuronal membrane potentials on the spatial pattern of n-sites.

In five simulations of spiking activity of the RefNN (Reference Neuronal Network) with the duration of 30 sec, the first one (top graph) was a control simulation with $V_i(t = 0) = V_{rest}$, and in each of four others the values $V_i(t = 0)$ were randomly taken from the uniform distribution in the interval between the resting potential $V_{rest}$ and threshold $V_{th}$.

All five graphs have the same structure as that in Fig. S2: At left, network spiking activity chunked into three vertically stacked graphs for 10 sec each. At right, schematic reconstruction of the corresponding spatial pattern of primary n-sites. The n-sites from the control simulation are shown by filled gray circles in all graphs as a landmark and the simulation-specific n-sites are colored in violet, with color saturation proportional to the relative activity of the corresponding n-site.

From the data presented, it can be seen that the diversity of the initial potentials leads to a redistribution of the relative activities of previously detected n-sites, but not to the emergence of new ones. The activity change is more pronounced for the n-sites, which are relatively faint in the benchmark (“control”) ReNN case, where all $V_i(t = 0)$ are equal.
