## Supplementary Figure S4 for "The vitals for steady nucleation maps of spontaneous spiking coherence in autonomous two-dimensional neuronal networks"

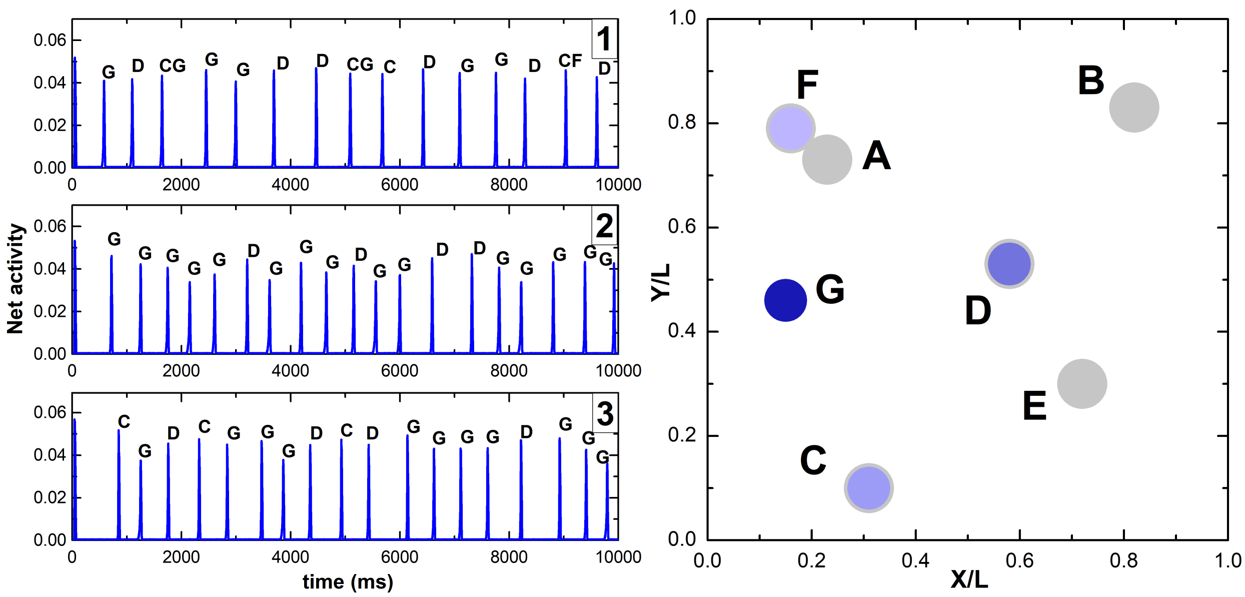


**Figure S4**. Influence of simulation time step $dt$ (see Sec. 2.3 in the main text) on the spatial pattern of n-sites. In three simulations of spiking activity of the RefNN (Reference Neuronal Network) with the duration of 10 sec, the first one (top left graph, labeled by "1" in its top right corner) was with $dt = 0.01$ ms (instead of $dt = 0.1$ ms as in all Figures shown in the main text) and the time bin $\triangle t = 0.2$ ms (instead of $\triangle t = 2$ ms) for averaging the network activity defined by Eq. (17) in the main text. That is, both $dt$ and $\triangle t$ were decreased tenfold. In the second simulation (middle left graph "2"), along with the above decrease of $dt$ and $\triangle t$, we also decreased tenfold the minimal spike-propagation delay $\tau_{del,min}$ (see Sec. 2.1.1, at the end), setting it $\tau_{del,min} = 0.02$ ms instead of standard 0.2 ms. Finally, in the third simulation (bottom left graph "3"), in addition to all changes made in the second one, we increased tenfold the spike propagation speed $v_{sp}$, setting it $v_{sp} = 2$ mm/ms instead of standard 0.2 mm/ms.

The right graph is a schematic reconstruction of the corresponding spatial patterns of primary n-sites for all three simulations. The n-sites from the standard RefNN simulation are shown by filled gray circles as a landmark and the simulation-specific sites are colored in violet, with color saturation proportional to the relative activity of the corresponding n-site. These data show that changing the time step $dt$ drastically modifies the spatial nucleation pattern: most n-sites (A, B, E) have vanished, and one new and very active n-site (G) has occurred. Simple rescalings of $\tau_{del,min}$ and $v_{sp}$ do not affect this result essentially (see Sec. 3.5 in the main text).
